# Adult dorsal root ganglion glial cells exhibit neurogenic potential upon reprogramming

**DOI:** 10.1101/2024.04.09.588701

**Authors:** Annemarie Sodmann, Niels Köhler, Nastaran M. Esfahani, Nina Schukraft, Annemarie Aue, Sara E. Jager, Maximilian Koch, Thorsten Bischler, Fabian Imdahl, Tom Gräfenhan, Enrico Leipold, Heike L. Rittner, Robert Blum

**Author notes:** **Corresponding author:**Prof. Dr. Robert Blum. RB and HLR share the senior author position.

## Abstract

In dorsal root ganglia (DRG), neuronal loss has been reported in patients with neuropathic pain, raising the question of whether the DRG, as part of the peripheral nervous system (PNS), harbor an endogenous cell source for neural repair. We found that adult mouse DRG harbor glial cells that dedifferentiate *in vitro* into Sox2/Sox10-positive glial progenitor-like cells. Coexpression of the developmental transcription factors Neurog1 and Neurog2 was sufficient to induce both neuronal and glial phenotypes. Nerve growth factor supported the maturation of a subset of neurons into nociceptor-like cells expressing functional TrpA1, TrpV1, and TTX-resistant Na_V_ channels. We report the limitation that we miss factors allowing consistent maturation to the sensory neuron profile. In summary, in the PNS, adult DRG-derived glial cells can acquire neural progenitor-like properties, show bipotent reprogramming competence, and may serve as an intrinsic cell source for sensory circuit regeneration.

## Introduction

Neuropathic pain following peripheral nerve injury severely impairs patients’ quality of life.^1,2^ Its underlying mechanisms are diverse, involve multiple cell types, and may differ between sexes.^3–5^ They include sensory neuron dysfunction following nerve injury or during the reinnervation of affected body regions,^6^ ectopic activity of sensory neurons in the dorsal root ganglia (DRG) or dorsal horn,^7,8^ aberrant axonal regeneration and activity from non-avulsed nerve roots,^9^ and the loss of peripheral sensory neurons or neural tissue within the DRG.^10–12^ For instance, patients with plexus injury, e.g. after motorcycle accidents, suffer from lifelong flaccid paralysis, sensory loss, and intractable pain. About half of these patients exhibit complete loss of the neural tissue in the injury-affected DRG.^13^ These observations raise the question of whether restoring sensory neurons together with their supportive glial environment, including satellite glial cells (SGCs), could promote sensory recovery^14^ and, in selected cases, alleviate neuropathic pain. Intriguingly, neonates with plexus injury show substantial sensory recovery and rarely develop neuropathic pain,^15^ suggesting that developmental or age-dependent regenerative mechanisms may influence the outcome following severe peripheral nerve injury.

In the central nervous system (CNS), harnessing endogenous glial cells *in situ* represents a promising strategy for brain repair at the site of injury.^16^ This approach may circumvent some of the challenges associated with allogeneic cell transplantation for neural tissue repair, including immune rejection, limited maturation of transplanted cells, and insufficient functional integration into neural circuits.^17^ In the CNS, glial cells can retain latent neural progenitor-like properties and, under specific experimental conditions, can be reprogrammed toward neuronal fates.^18,19^ In the peripheral nervous system (PNS), however, comparable neural progenitor-like potential has not been established. Such plasticity has been proposed for SGCs,^20,21^ whereas Schwann cells have not been shown to retain comparable latent progenitor-like potential.^22^

Following injury in the CNS, astrocytes acquire several hallmarks of neural progenitor cells, accompanied by increased expression of glial fibrillary acidic protein (GFAP).^23,24^ In the PNS, neuronal stress or peripheral nerve injury similarly induces a pronounced increase in GFAP expression in SGCs in close contact with sensory neurons.^14,25,26^ GFAP is also robustly expressed in Nageotte nodules, the non-neuronal cell clusters that form after sensory neuron death.^27^ These injury-induced phenotypic changes indicate increased cellular plasticity and may reflect a shift toward a more immature glial state.^23^ Pluripotent and progenitor-like cells have been identified in both embryonic and adult DRG^28–30^ and may differentiate into neurons after injury.^31–34^ Moreover, i*n vitro*, SGCs exhibit features reminiscent of Schwann cell precursors (SCPs),^35,36^ and others studies have reported differentiation of early postnatal rat DRG glial cells into neurons following treatment with small molecule inhibitors.^37,38^ Together, these observations raise the possibility that adult DRG glia may retain a latent progenitor-like state with the capacity for neuronal differentiation.

Human sensory neurons are commonly generated from induced pluripotent stem cells (iPSCs) using small-molecule inhibitors.^39,40^ However, these iPSC-derived sensory neurons show considerable heterogeneity in their transcriptional profiles between differentiations,^41^ and iPSC-derived populations do not yet fully recapitulate the molecular diversity of native DRG sensory neurons.^42^ An alternative approach uses forced expression of transcription factors, e.g., combinations of Neurog1, Neurog2 (neurogenins), or Brn3a, to induce sensory neuron phenotypes from iPSCs.^43,44^ This approach may provide more direct control over neuronal fate specification.^45^

The outcome of direct neuronal conversion strategies depends not only on the transcription factors expressed but also on the identity and developmental state of the starting population. For instance, neurogenins are used to induce sensory neurons from fibroblasts,^46–48^ and can also promote the generation of cortical-like glutamatergic neurons from human pluripotent stem cells, astrocytes, or other glial cells.^19,49,50^ In a reprogramming approach toward human nociceptors, differentiation proceeded through a SOX10+ neural crest intermediate stage,^51^ highlighting the neural crest lineage as a productive entry point for sensory neuron generation.

During development, neural crest cells give rise to sensory neurons in two waves of neurogenesis driven by distinct transcriptional programs.^52,53^ The first wave is mainly associated with Neurog2, whereas the second is dominated by the expression of Neurog1, although the two programs partially overlap.^54^ At later developmental stages, Brn3a and Islet1 contribute to sensory neuron differentiation, including the activation of transcription factors of the Runx family.^55,56^ Runx1, together with signaling through the nerve growth factor receptor TrkA, controls the specification of peptidergic and non-peptidergic nociceptors.^57–61^ In addition, the transcription factor Prdm12 contributes to nociceptive neuron development by regulating TrkA expression.^62,63^ These developmental programs, particularly the activity of the neurogenins, provide a framework for testing whether adult DRG glial cells retain the capacity to be reprogrammed toward sensory neuronal fates.

In this study, we show that SGC-like cells isolated from adult DRGs acquire a rapidly proliferative, progenitor-like state *in vitro*, suggesting that their removal from the sensory neuron niche induces a more immature cellular state. In this state, they retain bipotent reprogramming competence and can be induced to generate sensory neuron-like cells as well as glial cells following expression of neurogenins. Our findings provide evidence for retained developmental plasticity in the adult PNS.

## Results

### Satellite glial cells become fast-dividing glial progenitor-like cells *in vitro*

In the DRG cell culture, obtained from 4-8 weeks old wt C57BL/6 mice, we observed fast-dividing cells growing around DRG neurons. These cells were not visible immediately after plating but appear around single DRG neurons within a few days (Fig. 1 A). Immunofluorescence staining indicated that these cells originated from Fabp7/Sox2-positive SGCs tightly wrapped around the DRG neurons (Fig. 1 B). Already one day after plating, SGCs began losing contact with sensory neurons and became less Fabp7-positive, such that by day *in vitro* (DIV) 3 and 7, only a few Sox2-positive cells remained Fabp7-positive (Fig. 1 B-C). At DIV 7, cells abundantly expressed the sensory progenitor and glial cell markers Sox2 and Sox10 (Fig. 1 C). In addition, the culture contained smaller populations of NG2/PDGFRβ-positive cells (glial cells or pericytes), a few fibronectin-positive cells, possibly fibroblasts, and S100β- or GFAP-positive glial cells. To better characterize the cultured cells, we determined the mRNA-transcriptome of six independent DRG cultures, which showed homogenous transcripts per million (TPM) gene count distribution across cultures (Fig. S1 A, Table S1). Marker genes were chosen based on a published scSeq dataset of acutely dissociated DRG.^35^ The cultures expressed marker genes of multiple DRG cell types, including SGCs, Schwann cells, endothelial cells, fibroblasts, macrophages, myeloid cells, neurons, and pericytes, and closely resembled other cultured mouse DRG preparations (Fig. 1 D, Fig. S1 B). Some marker genes overlapped between cell types; for example, *Apoe* was abundantly expressed and attributed to both SGCs and macrophages. Generally, cells abundantly expressed marker genes for SGCs, such as *Cdh19* and *Sox2*. Furthermore, in line with the immunofluorescence staining, they also showed expression of the glial markers *Fabp7*, *Glul,* and *Sox10*, the pericyte marker *Pdgfrb*, and fibronectin (*Fn1*) (Fig. 1 E). We therefore concluded that the culture mainly contained peripheral glial cells, alongside smaller populations of other DRG cell types such as fibroblasts, pericytes, and macrophages.

**Fig. 1.**
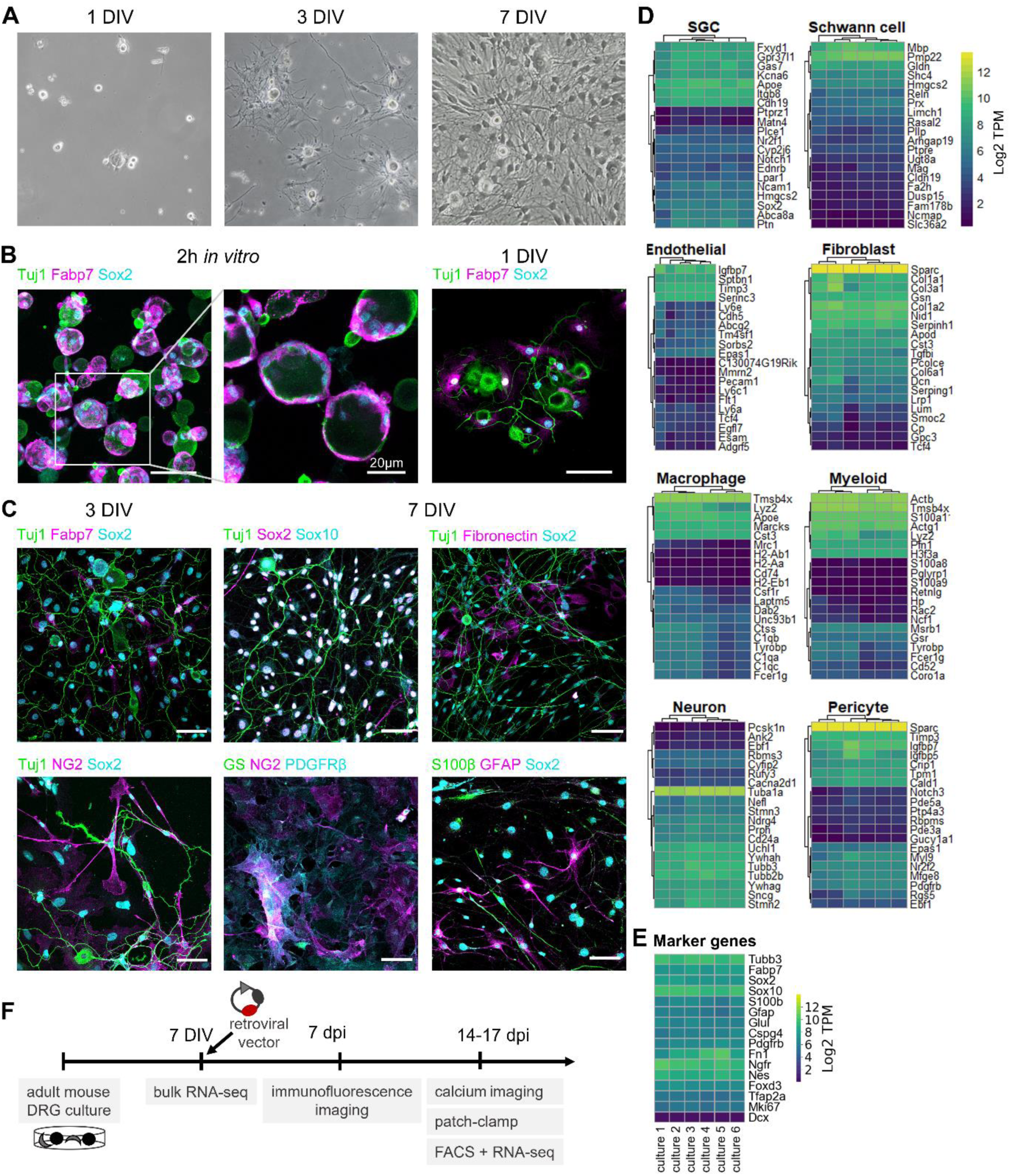
Fast-dividing glial progenitor-like cells in DRG cell cultures from adult mice. (A) Representative brightfield images of mouse cell culture at d 1, d 3, and d 7 *in vitro*. (B) Immunofluorescence images of DRG cell cultures at 2h and day *in vitro* (DIV) 1 labeled with the neuronal marker Tuj1 (βIII-Tubulin), the satellite glial cell (SGC) marker Fabp7, and Sox2, a progenitor and glial marker. (C) At DIV 3 and 7, DRG cultures were additionally stained for the glial and pericyte markers NG2 and PDGFRβ, the glial markers GFAP, GS, and S100β, the progenitor and glial cell marker Sox10, and the fibroblast marker fibronectin. Scale bars: 50 µm. (D) Heatmaps visualize the expression profile of marker genes of 8 cell types (SGC, Schwann cell (SC), endothelial, fibroblast, macrophage, myeloid, neuron, pericyte) across all samples. Marker genes were identified from publicly available scSeq data of acutely dissociated DRG.^35^ (E) Transcriptomic expression of marker genes of interest. Scale bar: Log2 transcripts per million (TPM). (F) Timeline of experimental procedures. DRG cells were prepared from adult mouse DRG and cultured for 7 days to perform RNA-seq or split and infected with a retroviral vector. Infected cells were analyzed with immunofluorescence imaging, patch-clamp, calcium imaging, and RNA-seq.

The high abundance of Sox2/Sox10-positive cells (Fig. 1 B, C), together with high mRNA expression of both markers (Fig. 1 E), further indicated a fast dividing, neural crest-derived glial cell type. Consistent with this, marker genes associated with neural crest cells, particularly those differentially expressed in boundary cap cells of the sensory neuron lineage,^54^ were also present (Fig. S1 C). Expression of these glial and progenitor marker genes was on the same level as housekeeping genes (Fig. S1 D). The transcriptome further confirmed abundant expression of markers typical of neural crest cells and neural progenitors, including Nestin (*Nes*), p75NTR (*Ngfr*), FoxD3 (*Foxd3)*, and AP2α (*Tfap2a)*, and the proliferation marker Ki67 (*Mki67*), whereas the immature neuron marker doublecortin (*Dcx*) was not expressed (Fig. 1 E).

Together, this shows that adult DRG glial cells can acquire a proliferative, progenitor-like phenotype characterized by expression of Sox2, Sox10, and neural crest-associated genes, *in vitro*.

### Retroviral vectors for bi- or tricistronic expression of transcription factors

The peripheral glial progenitor-like cells expressed Sox10 and Sox2, markers of the multipotent neural crest lineage from which sensory neurons and glial cells arise.^64,65^ We therefore hypothesized that these cells developed sufficient plasticity for transcription factor-mediated neuronal reprogramming. We selected the developmental transcription factors Neurog1, Neurog2, Brn3a (*Pou4f1*), Prdm12, and Runx1; all are key regulators of sensory neuron development and nociceptor specification and showed little endogenous expression in the cultured cells (Fig. S1 E). To ectopically express these factors, we generated a series of retroviral vectors. Starting with a previously established vector construct for glia-to-neuron reprogramming (Fig. S2 A),^19,50^ we cloned vectors expressing different combinations of Neurog1 (Ngn1), Neurog2 (Ngn2), Brn3a, Runx1, and Prdm12 (Table S2). In order to distinguish between the vector constructs in case of double infection, all vectors carried either an ^IRES^DsRed or ^IRES^GFP gene cassette (Zhao et al., 2006). We used codon-optimized transcription factor sequences, and included Flag, Myc, or HA affinity peptides in bi- or tricistronic expression constructs. The retroviral vector constructs were validated in HEK293 cells, which expressed each transcription factor (Fig. S2). In total, 15 different vector constructs were established.

### Adult peripheral glial progenitor-like cells have a high reprogramming potential

To test if the glial progenitor-like cells can be reprogrammed into neurons, mouse DRG cultures at DIV 7 were split and infected with different combinations of the retroviral vectors (Fig. 1 F). Within 7 days post infection (dpi), neurogenin-(Neurog1 or Neurog2) or Brn3a-expressing cells developed a neuronal Map2-/Tuj1-positive phenotype (Fig. 2 A). Bicistronic or tricistronic expression of Neurog1, Neurog2, and Brn3a induced neurons with even higher probability (Fig. 2 A). Computational quantification in tile-scans of whole coverslips showed that all ^IRES^DsRed-based vectors expressing neurogenin (Neurog1, Neurog2) and/or Brn3a induced significantly more neurons than a control vector expressing exclusively DsRed (Fig. 2 B-F). For example, 10-20% of cells infected with constructs encoding either Neurog1, Neurog2 or Brn3a expressed Tuj1 (Fig. 2 C) and about 5-10% were Map2 positive (Fig. 2 E). In contrast, expression of different combinations of these transcription factors increased the proportion of Tuj1-positive and Map2-positive cells to 40-50% and 20-30%, respectively (Fig. 2 C, E). Notably, the induction of neurons was most effective in cultures with tricistronic expression of all three factors (Brn3a-P2A-Ngn2-P2A-Ngn1) or with a bicistronic vector encoding both neurogenins (Ngn2-P2A-Ngn1). Other factors, including Runx1 and Prdm12, as well as their combination with other factors, showed no significant conversion potential (Fig. S3).

**Fig. 2.**
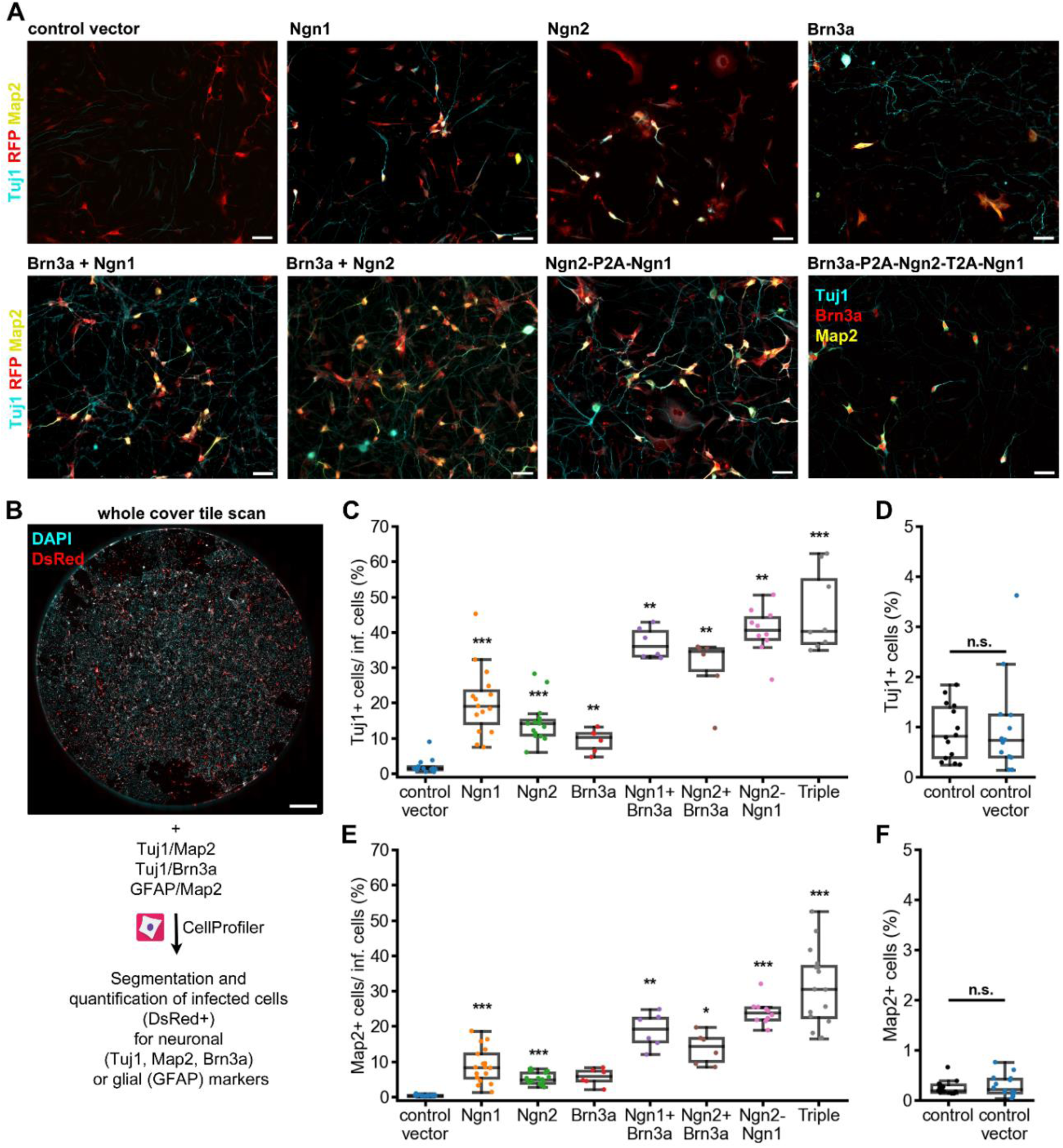
Reprogramming of peripheral glial progenitor-like cells with neurogenins and Brn3a. (A) Representative images of glial progenitor-like cells infected with CAG-IRES-DsRed-based retroviral vectors at 7 dpi. Infected cells are positive for DsRed (stained: RFP, red) or Brn3a (red). βIII-Tubulin (Tuj1, cyan) and Map2 (yellow) mark neuronal properties in cells transduced with retroviral vectors expressing indicated transcription factors. Brn3a and neurogenin vectors (Ngn1, Ngn2) were also simultaneously infected (Brn3a+Ngn1, Brn3a+Ngn2). Scale bars: 50 µm. (B) Whole cover tile scans were segmented and quantified with CellProfiler. Scale bar: 1 mm. (C-F) The percentage of the infected cells positive for Tuj1 (C, n=14/15/15/6/6/6/10/8; D, n=15/17/17/6/6/6/10/16) and Map2 (E, n=15/14; F, n=14/13) are shown with boxplots. The control vector was compared to an uninfected control based on all DAPI-positive cells. Triple: Brn3a-P2A-Ngn2-T2A-Ngn1. Groups of more than two were compared with the Kruskal-Wallis H-test and post hoc Mann-Whitney U test (C), or the Alexander-Govern test and post hoc Welch’s Test (E). Two groups were compared with the t-test (D), or Mann Whitney U test (F). Significant changes compared to the control vector are marked with: *p ≤ 0.05; **p ≤ 0.01; ***p ≤ 0.001; n.s.: not significant; with Bonferroni correction. Box plots show median and quartiles, with whiskers indicating 1.5× IQR and individual data points overlaid.

Because of the high reprogramming efficiency of Ngn2-P2A-Ngn1 (Fig. 2), we asked whether additional Brn3a expression, e.g., with the triple vector (Brn3a-P2A-Ngn2-T2A-Ngn1), was even necessary. Indeed, co-expression of Neurog1 and Neurog2 induced Brn3a protein expression in about 30% of the infected cells (Fig. 3 A-C). In uninfected or control infected cells, Brn3a immunoreactivity was barely detectable (Fig. 3 D). We concluded that forced co-expression of Brn3a along with both neurogenins can be omitted, so in subsequent experiments we focused on Ngn2-P2A-Ngn1 coexpression for reprogramming.

**Fig. 3.**
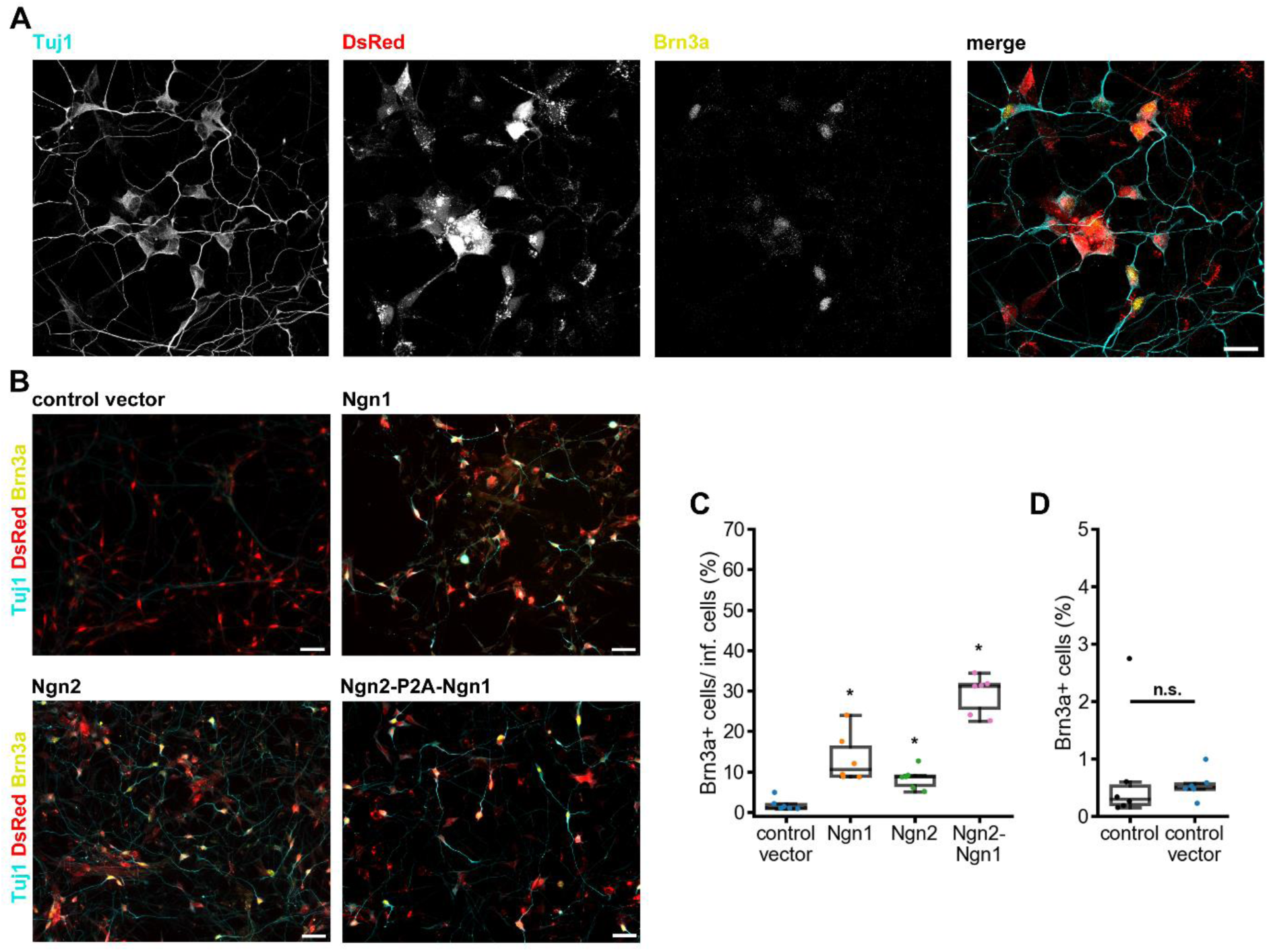
Brn3a expression in peripheral glial progenitor-like cells induced by neurogenins. (A-B) Representative fluorescent images of cells transduced with Ngn2-P2A-Ngn1 (A, confocal) or control- and neurogenin-transduced cells (B) at 7 dpi. Cells were labeled for the neuronal marker βIII-Tubulin (Tuj1, cyan) and the transcription factor Brn3a (yellow). ^IRES^DsRed marks infected cells. Scale bars: (A) 25 µm. (B) 50 µm. (C-D) Percentage of Brn3a-positive cells in neurogenin-infected (Neurog1 and/or Neurog2) cells (C, all n=6) or in the controls (D, uninfected or control vector infected, both n=6). Groups of more than two were compared with the Kruskal-Wallis H-test and post hoc Mann-Whitney U test (C); two groups were compared with the Mann Whitney U test (D). Asterisks indicate significant differences from the control vector: *p ≤ 0.05; n.s.: not significant; with Bonferroni correction. Box plots show median and quartiles, with whiskers indicating 1.5× IQR and individual data points overlaid.

### Neurogenin expression can also induce a GFAP-positive glial cell phenotype

Neurog2 has previously been shown to promote differentiation of peripheral neural crest-derived progenitor cells toward both sensory neuronal and glial fates during development, rather than restricting cells exclusively to the neuronal lineage.^66,67^ Consistent with these *in vivo* developmental studies, forced expression of both Neurog1 and Neurog2 was sufficient to also induce a GFAP-positive phenotype (Fig. 4 A-B), whereas co-expression of the proneuronal factor Brn3a suppressed this glial-like phenotype (Fig. 4 C). GFAP-positive cells were induced in about 20% of the neurogenin (Ngn1, Ngn2, Ngn2-P2A-Ngn1) infected cells (Fig. 4 D), compared to 0.5–1% in controls (Fig. 4 E).

**Fig. 4.**
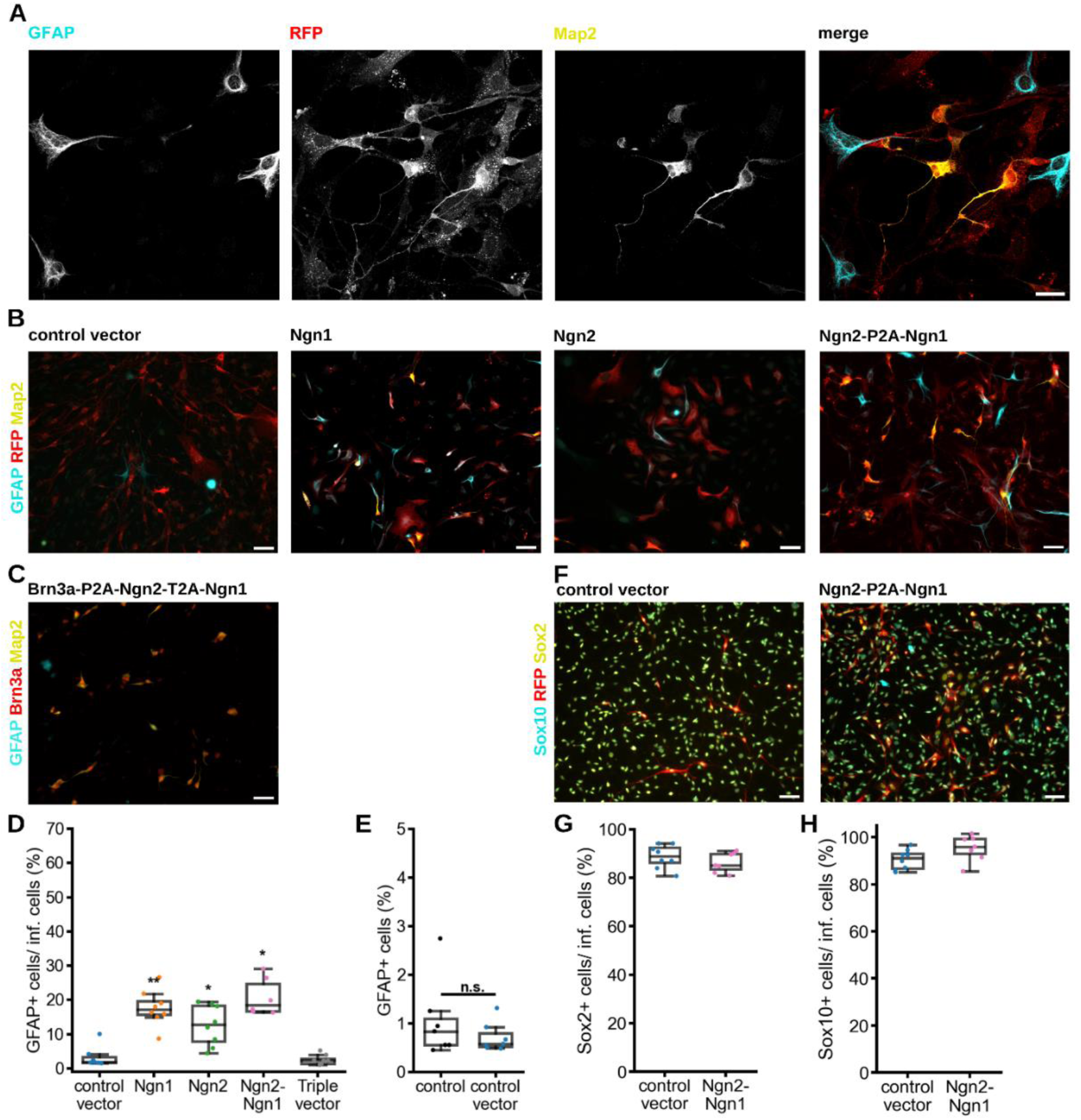
GFAP-positive glial cells induced by neurogenins. (A) Representative confocal image of Ngn2-P2A-Ngn1 infected cells (7 dpi) labelled for the glial marker GFAP (cyan), the neuronal marker Map2 (yellow), and RFP (red). Scale bar: 25 µm. (B) GFAP (cyan) and Map2 (yellow) in control and neurogenin-transduced (Neurog1 and/or Neurog2) cells. RFP (red) labels ^IRES^DsRed-positive, infected cells. Scale bars: 50 µm. (C) Representative staining of cells transduced with the triple vector Brn3a-P2A-Ngn2-T2A-Ngn1. Scale bar: 50 µm. (D-E) Percentage of GFAP-positive cells among neurogenin- or triple-vector-infected cells (D, n=8/7/8/8/6) and among controls (E, uninfected and infected, n=7). (F) Representative images of cells transduced with the control vector or Ngn2-P2A-Ngn1 (7 dpi), labelled for Sox10 (cyan), Sox2 (yellow) and RFP (red). Scale bars: 50 µm. (G-H) Quantification of tile-scans of individual cultures shows the percentage of Sox2-positive cells (G) or Sox10-positive cells (H) in control-(n=8) and neurogenin-infected (n=7) cells. Groups of more than two were compared with the Kruskal-Wallis H-test and post hoc Mann-Whitney U test (D); two groups were compared with the Mann Whitney U test (E), or t-test (G, H). Asterisks indicate significant differences from the control vector: *p ≤ 0.05; **p ≤ 0.01; n.s.: not significant; Bonferroni corrected. Box plots show median and quartiles, with whiskers indicating 1.5× IQR and individual data points overlaid.

Moreover, we characterized Sox2 and Sox10 expression in the transduced cell populations. Approximately 90% of cells in both the neurogenin and control groups expressed these markers (Fig. 4F–H), indicating that the majority of transduced cells exhibited a Sox2/Sox10-positive phenotype under the conditions examined. Although this finding is consistent with the presence of a peripheral glial progenitor-like state among the starting cells, it does not establish that the induced neuronal or GFAP-positive phenotypes arose specifically from Sox2/Sox10-positive cells. Definitive evidence for their cellular origin will require lineage-tracing approaches.

### Induced neurons show immunoreactivity to nociceptor markers

To further differentiate the induced neurons, 100 ng/ml NGF was added to the cell culture. NGF is a neurotrophin and high-affinity ligand for TrkA (tropomyosin receptor kinase A) that promotes the differentiation of sensory neurons.^61^ When cultured for more than two weeks (14-16 dpi), induced neurons developed longer neurites, had a diameter of about 10-15 µm, and were able to form clusters of neurons (Fig. 5 A-C). Moreover, cells showed immunoreactivity to marker proteins typical for nociceptors. Some of the neurons showed immunoreactivity for the intermediate neurofilament peripherin, TrkA, and the TTX-resistant voltage-gated sodium channel Na_V_1.9, which is typically expressed in nociceptive neurons of the DRG (Fig. 5 D-F).^68,69^ Also, induced neurons expressed the transient receptor potential channel TrpA1, the receptor for mustard oil, and TrpV1, the receptor for capsaicin (Fig. 5 G-H).

**Fig. 5.**
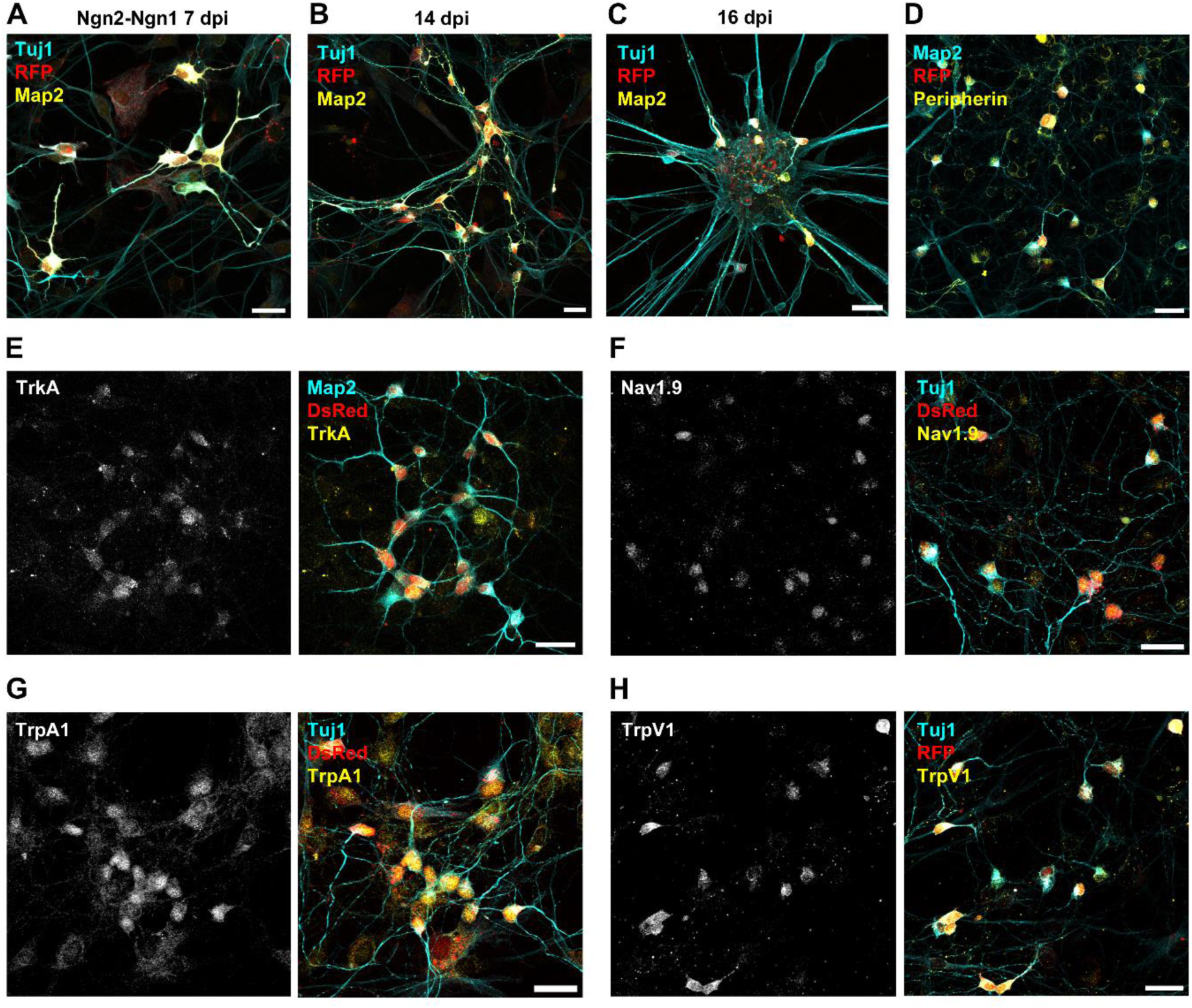
Induced neurons express markers characteristic for sensory/nociceptive neurons. Representative confocal images showing induced neurons reprogrammed with Ngn2-P2A-Ngn1, identified by expression of ^IRES^DsRed (stained: RFP, red). Scale bars: 25 µm. (A-C) Labelled are the neuronal markers βIII-Tubulin (Tuj1) and Map2 at 7 dpi (A), 14 dpi (B), and 16 dpi (C). (D-E) Cells labelled for Peripherin (D, yellow) or TrkA (E, yellow) together with Map2 (cyan) at 14 dpi. (F-H) Cells labelled for the sodium channel Na_V_1.9 (F, yellow), the nociceptor markers TrpA1 (G, yellow), or TrpV1 (H, yellow) counterstained with Tuj1 (cyan) at 14 dpi.

### Induced neurons are sensitive to capsaicin and mustard oil

To characterize induced neurons on a functional level, we performed calcium imaging and patch clamp recording. First, we tested induced neurons for calcium responses to the TrpV1 agonist capsaicin and the TrpA1 agonist mustard oil (allyl-isothiocyanate, AITC) at 14-15 dpi, as both ion channels mediate typical calcium responses in nociceptive DRG neurons harvested from adult animals.^70,71^ Neurogenin-infected cells were identified by their co-expression of DsRed, with uninfected cells on the same coverslip serving as controls (Fig. 6 A). Induced neurons responded to acute application of capsaicin (10 µM for 10 s) with a cytosolic calcium transient, heterogeneous in both timing and amplitude (Fig. 6 B), while uninfected cells showed little to no response (Fig. 6 C). Overall, about 35% of DsRed-positive cells were capsaicin-sensitive (Fig. 6 D). Stimulation of TrpA1 with AITC (100 µM, 10 s) induced calcium responses in 11% of the neurogenin-expressing, DsRed-positive cells (Fig. 6 E-G).

**Fig. 6.**
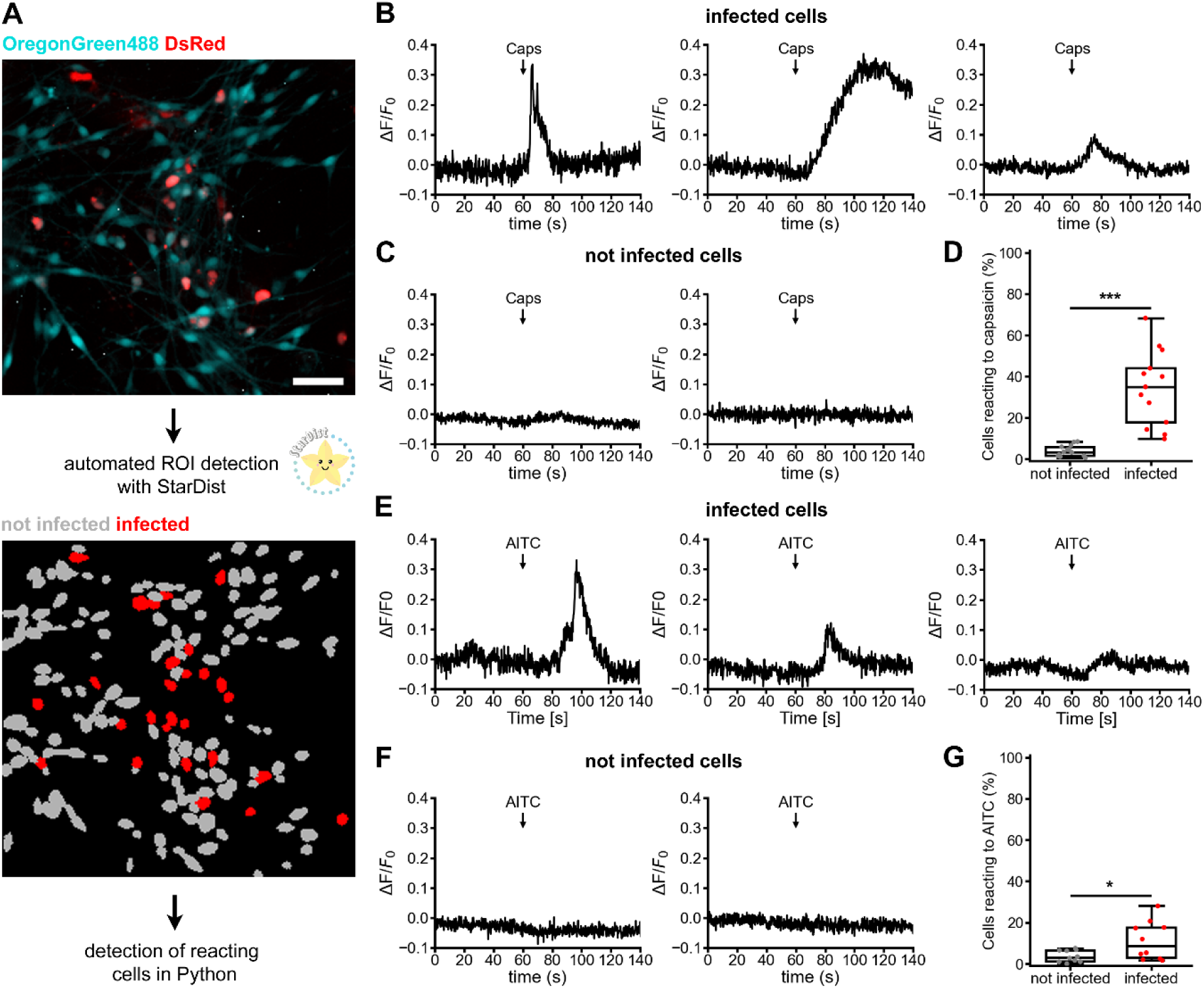
Sensitivity of induced neurons to the TrpV1 agonist capsaicin and the TrpA1-agonist AITC. (A) Workflow: Calcium signals were detected with Oregon Green 488 (cyan) in Ngn2-P2A-Ngn1 infected cells (14-16 dpi), labelled with ^IRES^DsRed (red). Automatic detection of the region of interest (ROI) with StarDist2D separated not infected (grey) from infected (red) cells. The percentage of reacting cells was determined using python scripts. Scale bar: 50 µm. (B-C) Representative calcium signals of infected (B) and uninfected (C) cells stimulated for 10 s with 10 µM capsaicin (Caps). (D) Box plot: Percentage of infected and uninfected capsaicin-sensitive neurons (n=13 measurements). (E–G) Representative AITC-induced calcium signals (E, F) and percentage of AITC-sensitive neurons (G) upon stimulation with 100 µM AITC for 10 s (n=10 measurements). Groups were compared with Welch’s t-test. Significant differences: *p ≤ 0.05; ***p ≤ 0.001. Box plots show median and quartiles, with whiskers indicating 1.5× IQR and individual data points overlaid.

### Electrophysiological properties of induced neurons

Next, we analyzed the electrophysiological properties of induced neurons at 14 to 17 dpi in whole-cell patch-clamp recordings (Fig. 7 A). Immediately after the whole-cell configuration was established, test pulses to -20 mV and 60 mV were applied to monitor the activity of voltage-gated ion channels. Using this stimulation paradigm, induced neurons gave rise to pronounced inward and outward currents, indicating expression of functional voltage-gated Na_V_ and K_V_ channels (Fig. 7 B, n = 72). Mean maximum inward current densities were smaller in induced (repro) neurons (-122.70 ± 12.04 pA/pF) than in naïve DRG neurons at DIV 1-2 (-337.38 ± 56.70 pA/pF) (Fig. 7 C). However, mean maximum outward current densities (repro: 497.07 ± 22.33 pA/pF; naïve: 525.37 ± 48.65 pA/pF) and resting membrane potential (repro: -46.95 ± 1.35 mV; naïve: -49.32 ± 2.07 mV) were indistinguishable between the two groups (Fig. 7 D, E).

**Fig. 7.**
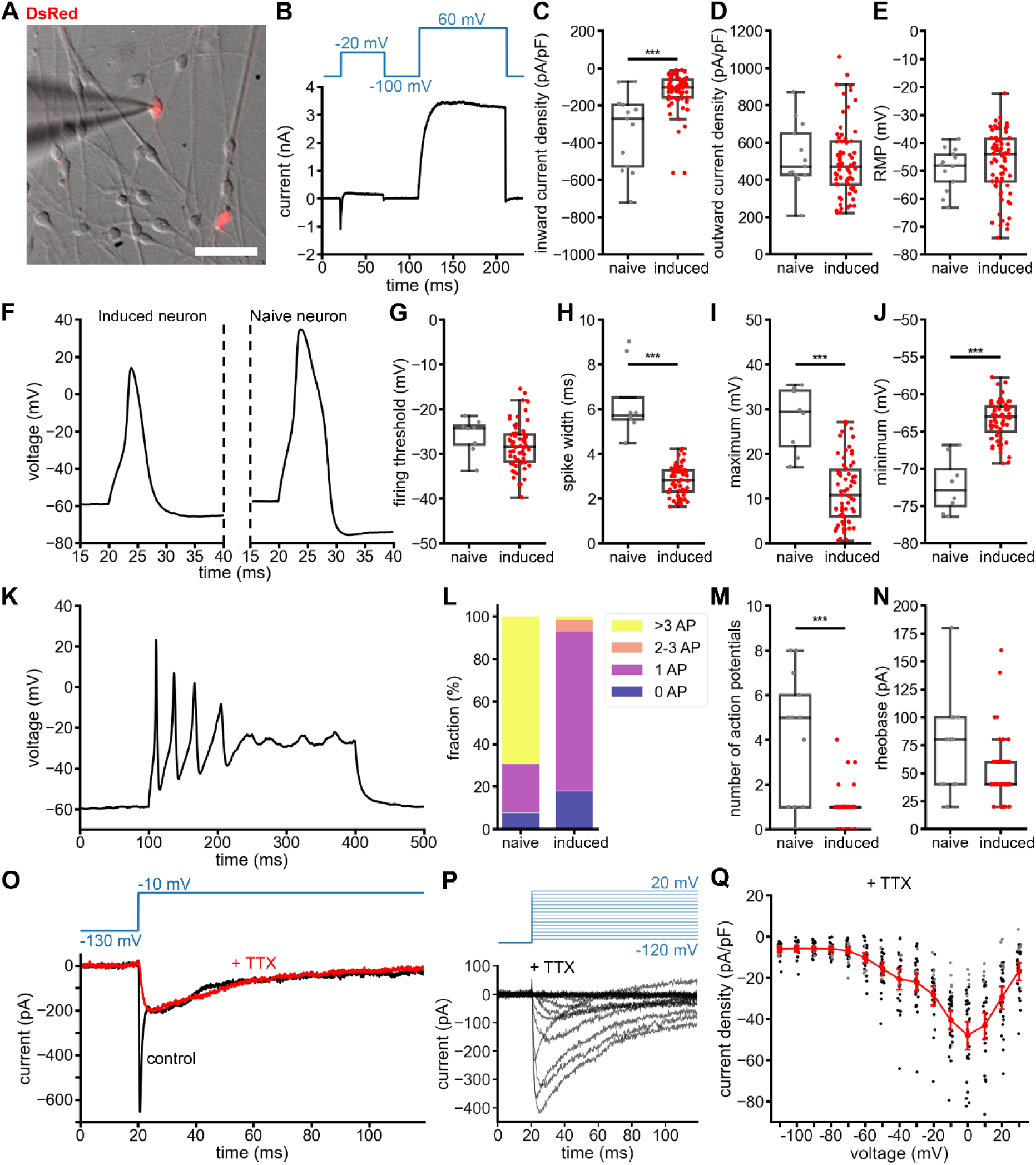
Electrophysiological properties of induced neurons. (A) Microscope image of induced neurons at 17 dpi, obtained during a patch-clamp recording session. Cells expressing the infection marker DsRed were identified by their bright red fluorescence. Scale bar: 50 µm. (B) Representative currents of an induced neuron at -20 and 60 mV. (C-E) Comparison of mean maximum inward (C) and outward (D) current density, and resting membrane potential (RMP) (E) of induced neurons (14-17 dpi, n=72) to those of naïve small-diameter DRG neurons at DIV 1-2 (n=13). (F) Representative single action potentials (APs) of an induced neuron and naïve DRG neuron stimulated for 3 ms with 60 pA or 220 pA, respectively. (G-J) Firing threshold (G), spike width at -20 mV (H), maximum voltages (I), and minimum voltages (J) of single action potentials of naïve and induced neurons. (K) Induced neuron firing multiple action potentials in response to a 300 ms current injection of 40 pA. (L) Fractions of naïve and induced neurons firing a maximum of zero, one, two or three, or more action potentials during a series of current injections. (M-N) Number of action potentials (M) and rheobase (N) of naïve and induced neurons. (O) Inward currents of a representative induced neuron at -10 mV before and after application of 500 nM TTX. (P) Representative current traces of an induced neuron in presence of 500 nM TTX in response to 100 ms long depolarizing voltage pulses ranging from -120 to 20 mV in steps of 10 mV. (Q) Mean peak inward current densities ± s.e.m. (red) as a function of voltage of induced neurons in the presence of 500 nm TTX. Dots represent peak current densities of individual cells that, at least once, exhibit a maximum inward current density greater (black, n=26 cells) or lower (gray, n=5 cells) than 30 pA/pF. Groups were compared with Mann Whitney U test (C-E, M), or t-test (G-J, N). Significant differences: ***p ≤ 0.001, Bonferroni corrected. Box plots show median and quartiles, with whiskers indicating 1.5 ×IQR and individual data points overlaid.

We next determined action potential characteristics in current-clamp mode. Brief depolarizing current injections evoked single action potentials in induced neurons, further supporting their neuronal phenotype (Fig. 7 F). However, the evoked responses differed significantly from those recorded in naïve DRG neurons. They were characterized by a take-off voltage of -28.18 ± 0.63 mV, a duration of 2.82 ± 0.08 ms (measured at -20 mV), and maximum and minimum peak voltages of 11.76 ± 0.91 mV and -63.44 ± 0.33 mV, respectively (Fig. 7 G-J). To evoke repetitive firing, cells were clamped to -60 mV and stimulated with escalating current injections (0 - 220 pA) for periods of 300 ms. Under this protocol, three or more overshooting action potentials (peak voltage > 0 mV) were triggered in the majority of naïve DRG neurons (69%), while most (75%) induced neurons fired only one action potential (Fig. 7 K-M). Only a small fraction (7%) of induced neurons fired repetitively, and even these showed rapidly decreasing amplitudes that ultimately failed to reach the 0 mV detection threshold (Fig. 7 K). Rheobase was comparable (repro: 51.19 ± 3.32 pA; naïve: 75.0 ± 12.58 pA) between induced and naïve neurons (Fig. 7 N).

Expression of TTX-resistant (TTXr) voltage-gated sodium channels (Na_V_1.8, Na_V_1.9) is a hallmark feature of nociceptive neurons (Dib-Hajj et al., 2002, Djouhri et al., 2003) and their role in peripheral nociception is well documented.^71,72^ To test for functional expression of TTXr Na_V_ channels, induced neurons were analyzed in a separate set of whole-cell voltage-clamp recordings using optimized buffers supplemented with 500 nM TTX to specifically isolate TTXr current components.^73^ While the TTXs current component of induced neurons inactivated rapidly within a few milliseconds, the TTXr current component inactivated considerably slower (Fig. 7 O). Induced neurons generated TTXr Na^+^ inward currents of up to about 400 pA in response to a 0 mV test pulse (Fig. 7 P). Using a current density of 30 pA/pF as the detection limit, TTXr Na^+^ currents were observed in 26 of 31 induced neurons tested (84%, Fig. 7 Q). Moreover, the peak current densities of TTXr currents showed a biphasic voltage-dependence, with one component peaking around -40 mV and a second component with a minimum at 0 mV, potentially reflecting the voltage-dependent activation of Na_V_1.9 and Na_V_1.8, respectively.^74^

In summary, the electrophysiological data show that the induced neurons are electrically excitable, though less than naïve DRG neurons. Consistent with our immunofluorescence labelling (Fig. 5), a substantial proportion of the induced neurons may express functional TTXr Na_V_ channels.

### Variability of reprogramming outcome

Building on these findings, we next assessed the robustness of the reprogramming protocol across experimental batches and observed variability in both the number and phenotypic identity of induced neurons (Fig. S4). After three years of experiments, a laboratory relocation coincided with a shift in the characteristics of the reprogrammed cells (Fig. S5 A). We found that batch-to-batch variation in growth factors and serum was linked to differences in neuronal induction and Brn3a expression in the Neurog2-Neurog1-DsRed-infected cells (Fig. S4 B,C). We therefore standardized serum sourcing and supplemented cultures with additional neurotrophic factors (BDNF, GDNF) to improve consistency. While this stabilized several aspects of the protocol, the full nociceptor-like phenotype observed in earlier experiments, including immunoreactivity for TRPV1 and TRPA1 and capsaicin responsiveness, could not be consistently reproduced under these conditions (Fig. S4 D–F). Importantly, however, neurogenin-infected cells consistently displayed Map2 and βIII-tubulin immunoreactivity, and responded robustly to depolarization with 90 mM KCl (Fig. S4 F). This suggests that basic neuronal conversion remained achievable, even as the specific nociceptor-like sub-phenotype proved more sensitive to culture conditions.

### Single cell RNA sequencing of neurogenin-infected cells shows generation of immature neurons

Next, we wanted to resolve the transcriptomic identity of the Neurog2-Neurog1-DsRed-infected cells more precisely. To this end, we performed single cell RNA sequencing (scRNA-seq) of FACS-sorted DsRed-positive cells at 14 dpi, cultured in the presence of 100 ng/ml NGF (Fig. 8 A). Immunofluorescence staining confirmed that sorted cells were DsRed and Tuj1 positive three days after sorting. Following 10x Genomics’ scRNA-seq, data from 4,549 individual cells were analyzed for cell type and marker gene expression. Cells were divided into six cell clusters (Fig. 8 B). Based on established marker genes, five of the six clusters were identified as glial progenitors (two clusters, expression of Sox2, Sox10, Fabp7, Cdh19), immature neurons (Map2, Dcx), fibroblasts (Fn1), and macrophages (Aif1) (Fig. 8 C). The remaining “Unidentified” cluster expressed high levels of ribosomal RNA and genes such as Gm42418 and AY036118 (Fig. 8 D), indicating an rRNA contamination phenotype^75^ likely representing a technical artifact^76^ caused by ambient RNA contamination, so it was excluded from further analysis.

**Fig. 8.**
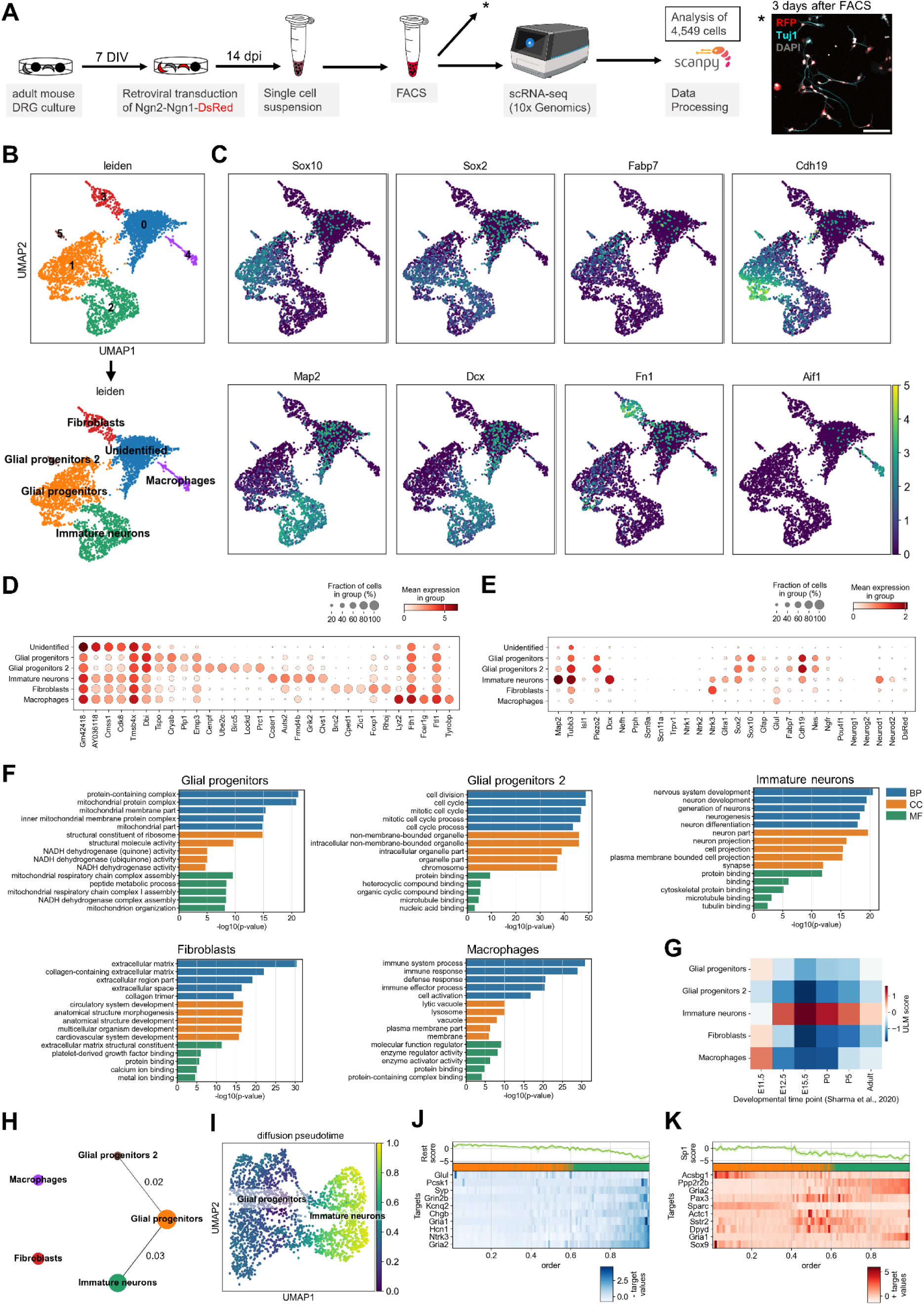
scRNA sequencing of neurogenin tranduced cells shows an immature neuronal phenotype. (A) Neurog1-Neurog2-DsRed transduced cells were cultured for 14 days, FACS sorted for DsRed, and 10x Genomics’ single cell RNA-sequencing (scRNA-seq) was performed. FACS sorted cells, cultured, and stained after 3 days, were RFP and βIII-Tubulin (Tuj1) positive. scRNA-seq data from 4,549 cells were processed using scanpy. (B) Leiden clustering of cells into six clusters (Unidentified, Glial progenitors, Immature neurons, Fibroblasts, Macrophages, Glial progenitors 2). (C) Expression of marker genes for progenitor cells (Sox2, Sox10), satellite glial cells (Fabp7, Cdh19), immature neurons (Map2, Dcx), fibroblasts (Fn1), and macrophages (Aif1) in single cells, visualized in UMAPs. Color intensity represents normalized gene expression (0–5). (D-E) Mean expression of highly differentially expressed marker genes (D), and of additional genes of interest for neurons, nociceptors, glial progenitors, and retroviral vector expression (E) in the six clusters. Dot size represents the proportion of cells within each cluster expressing the respective gene, while color intensity indicates the mean expression level. (F) Gene ontology (GO) terms associated with the top 200 differentially expressed genes. Shown are the top 5 terms of biological processes (BP, blue), cellular component (CC, orange), and molecular function (MF, green). (G) Enrichment analysis of genes enriched across developmental time points from E11.5 to adulthood in DRG sensory neurons.^61^ Scale bar: ULM score (mean). (H) Partition-based graph abstraction (PAGA) graph of the cell clusters. (I) UMAP of the diffusion pseudotime after PAGA of the “Glial progenitors” to the “Immature neurons” cluster. (J-K) Pseudotime enrichment analysis of regulated transcription factors and their targets using the CollecTRI network. Mean expression of the two highest scored transcription factors, *Rest* (J) and *Sp1* (K), and their negative (blue) or positive (red) targets are visualized along the pseudotime trajectory order.

The neuronal cluster displayed an immature phenotype, characterized by expression of Map2, Tubb3, Dcx, Sox2, and the neuronal bHLH transcription factors Neurod1 and Neurod2, which are expressed downstream of *Neurog1* and *Neurog2* during sensory neurogenesis.^54^ In contrast, mature nociceptor marker genes, including *Nefh* (neurofilament heavy chain), *Prph* (peripherin), *Scn9a* (Na_V_1.7), *Scn11a* (Na_V_1.9), *Trpv1* (TrpV1), *Ntrk1* (TrkA), and *Ntrk2* (TrkB), were not detected in the sorted cells (Fig. 8 E). Instead, the GDNF receptor *Gfra1* and *Ntrk3* (TrkC) were expressed in the neuronal cluster, which are both important for nervous system development and neuronal survival.^77,78^ The glial progenitors expressed high levels of *Cdh19*, *Plp1*, *Nes*, *Sox2*, and *Sox10*, and lower levels of *Fabp7, Ngfr*, and *Gfap*. Gene ontology (GO) analysis of genes enriched in the two glial progenitor clusters distinguished them by signatures of mitochondrial protein complex and NADH dehydrogenase activity terms or increased mitotic activity, respectively. The fibroblasts were associated with extracellular matrix, and macrophages with immune system processes. The immature neurons were associated with GO terms related to nervous system development, neurogenesis, neuron differentiation, and neuron part (Fig. 8 F). Moreover, they exhibited the strongest association with the E15.5 sensory neuron developmental stage characterized by Sharma et al.^79^ (Fig. 8 G).

Partition-based graph abstraction (PAGA) identified connections between the glial progenitor cluster and both the immature neuron cluster (connectivity = 0.03) and glial progenitor 2 cluster (connectivity = 0.02), whereas no connectivity was detected between the remaining clusters (excluding the unidentified cluster) (Fig. 8 H). We further characterized the transcriptional transition from glial progenitor to immature neurons. The embedding shows a continuum of intermediate cell states bridging the populations (Fig. 8 B). Therefore, we performed a diffusion pseudotime (DPT) analysis of the two clusters, rooted in the glial progenitor cluster (Fig. 8 I). The most enriched transcription factors that declined along the pseudotime trajectory were *Rest* (Fig. 8 J), a repressor of neuronal genes^80^ and competitor of Neurog2,^81^ and *Sp1* (Fig. 8 K), whose expression declines during neuronal differentiation.^82^ Negative targets of *Rest* showed an increase in expression along pseudotime, for example the glutamate receptor subunits *Gria1* and *Gria2*, the neurotrophic receptor tyrosine kinase 3 (*Ntrk3*), and the synaptic vesicle protein synaptophysin (*Syp*). Positive targets of *Sp1* showed different gene expression patterns. *Sparc* and *Sox9* decreased over pseudotime, whereas *Pax3*, *Actc1*, and *Sstr2* showed peak expression at intermediate pseudotime order.

Together, these transcriptomic analyses support the conclusion that Neurog2/Neurog1 expression induces the generation of immature neurons from glial progenitors, as evidenced by the transcriptional continuum and pseudotime trajectory connecting both populations. This confirms that adult DRG-derived glial cells exhibit neurogenic potential.

### High similarity to human DRG cell cultures

To determine whether these findings might be transferable to human DRGs, we investigated whether peripheral glial progenitor-like cells are also present in human DRG cultures. To this end, we compared our bulk RNA-seq dataset from mouse DRG cultures at DIV 7 with a publicly available dataset from human DRG cultures at DIV 8.^83^ The 1,000 most highly expressed genes in both datasets were associated with terms such as gliogenesis, cell adhesion, and nervous system development (Fig. 9 A). Of these, 399 genes were shared between species, and the enriched biological processes showed substantial overlap (Fig. 9 B). We identified 15,476 one-to-one orthologues between the mouse and human datasets, which exhibited a strong correlation in z-scored mean expression (Spearman’s ρ = 0.823; Fig. 9 C). Marker genes of interest, including *SOX10*, *SOX2*, *GLUL*, *FN1*, *NGFR*, and *NES* were robustly expressed in both species, with comparable expression levels (Fig. 9 D). Only *TUBB3* showed lower expression in human DRG cultures. We also identified species-biased gene expression. Human-biased genes included *RN7SL1* (RNA component of signal recognition particle), *CRYGC* (lens crystallin), and *H2BC18* (histone), whereas *EEF1G* (translation elongation factor), *FXYD1* (ion transport regulator), and *HMGCS2* (ketogenesis^84^) were more abundant in the mouse data (Fig. 9 C,E). Gene Ontology analysis of the top 300 human-biased genes identified enrichment for biological processes related to L-tryptophan/indole metabolism, muscle cells, and GPCR-signaling (Fig. 9 F). For mouse, we found higher regulation of Wnt signaling pathways, sodium ion transport by FXYD-family genes, and nervous system development (Fig. 9 G).

**Fig. 9.**
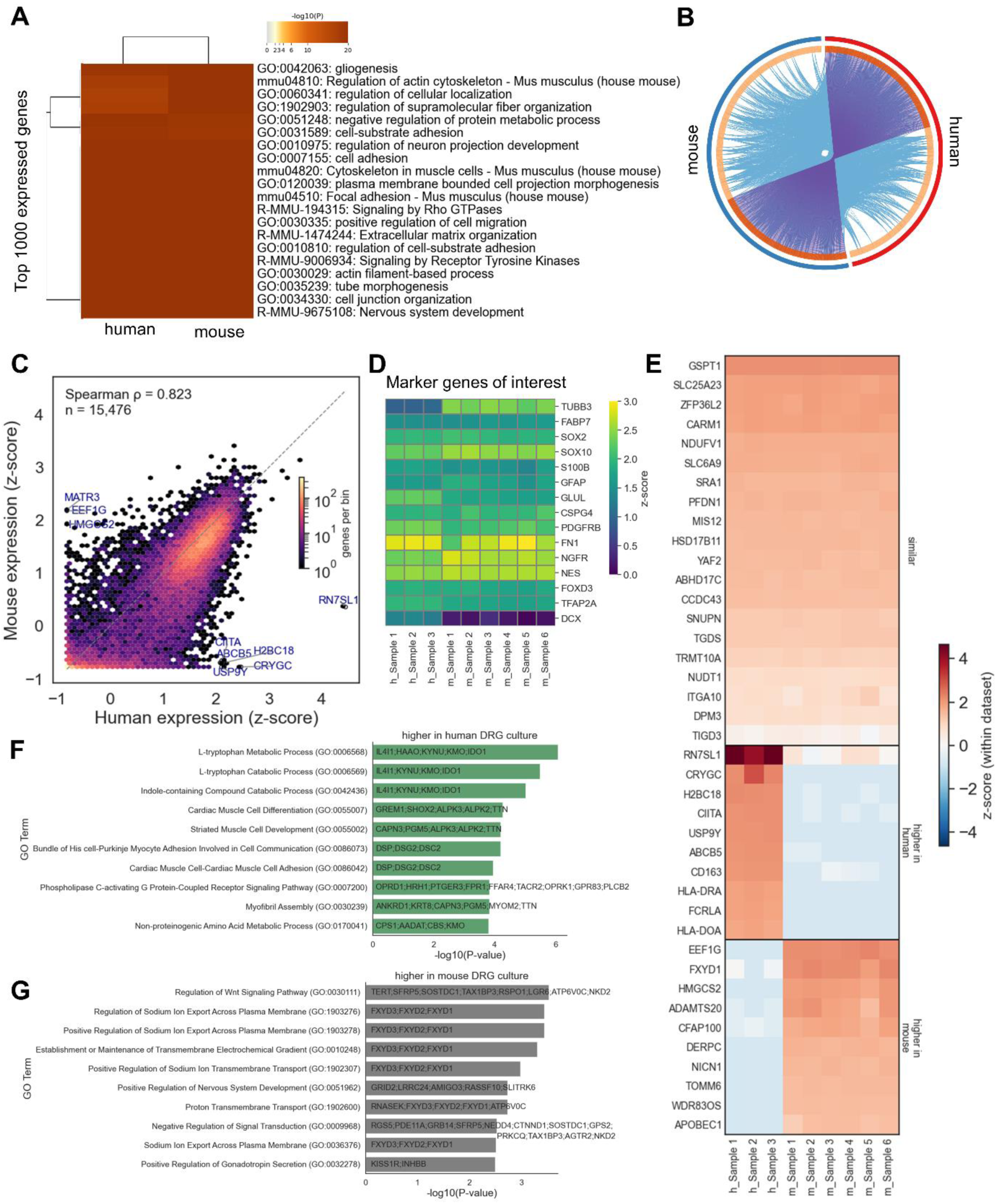
High similarity between mouse and human DRG cell culture. (A) Heatmap of enriched terms associated with the top 1000 expressed genes in the mouse (DIV 7) or human^83^ (DIV 8) DRG culture dataset. (B) Circos plots of overlaps between these top 1000 genes. Purple lines link identical genes between conditions, and blue lines link genes with shared ontology terms. (C) Cross-species concordance of DRG culture transcriptomes. Per-gene expression (mean z-scored within each dataset) for human versus mouse across n = 15,476 one-to-one orthologues. Hexagonal bins show the density of all 1:1 orthologues (color = genes per bin, log scale); individually labelled points mark the genes with the largest off-diagonal residuals. Overall rank concordance was assessed by Spearman’s rank correlation (ρ = 0.82). (D) Expression of marker genes of interest. Scale: Z-score within each dataset. (E) Concordant and species-biased gene expression between human and mouse DRG cultures. Within-dataset z-scored expression for human and mouse across selected 1:1 orthologues. Rows are grouped by the human–mouse residual (mouse z − human z): the 20 genes with the smallest absolute residual (similar), the 10 most human-biased (higher in human), and the 10 most mouse-biased (higher in mouse). Color encodes the z-score per species (red high, blue low). (F-G) Top 10 GO Biological Process terms associated with the top 300 genes higher in human (F) or higher in mouse (G), based on human–mouse residual. Visualized as bar plots ordered by −log₁₀(P-value) with contributing genes annotated above each bar.

Together, these findings demonstrate that mouse and human DRG cultures share a similar transcriptional landscape, including expression of key peripheral glial progenitor markers, while exhibiting a limited number of differences that may reflect species-specific biology or differences in culture conditions.

## Discussion

This study shows that SGC-like cells from adult DRG have the genetic plasticity to dedifferentiate into glial progenitor-like cells *in vitro*. Retroviral co-expression of the early fate determinants for peripheral neural cells, Neurog1 and Neurog2, was sufficient to induce neurons with properties of sensory neurons *in vitro*. The peripheral neural cell source shows at least bipotency, as neurogenin-mediated reprogramming could also induce a GFAP-positive glial cell fraction.

The peripheral glial progenitor-like cells expressed Sox2 and Sox10, and markers for SGCs (e.g. *Cdh19*, *Fabp7*), and boundary cap cells. We suspect that they originate from SGCs tightly attached to DRG neurons,^85,86^ because cell growth started around single, cultured sensory neurons (Figure 1 A-B). Typical tissue dissection protocols cannot fully separate the SGC envelope from isolated DRG neurons, suggesting that these cells, when taken out of their niche, adhere to the 2D cell culture surface, and start to proliferate. In this state, cells rejuvenate and express features similar to sensory progenitor cells (e.g. Sox2, Sox10). A previous study suggested that SGCs represent precursor cells in the Schwann cell lineage, because they show transcriptional and morphological similarities to Schwann cells, and are able to myelinate purified sensory neurons *in vitro.*^36^ However, SGCs *in vitro* differ from cells characterized *in vivo*^35^ and have been shown to develop multipotent glial precursor properties.^87^ This study supports the view that SGCs in culture acquire properties of a neural crest-derived peripheral progenitor, with high developmental plasticity that extends beyond the Schwann cell lineage.

This plasticity potential might also be present in human DRGs, as cultured DRG from organ donors shared a similar transcriptomic profile with mouse DRG *in vitro*, characterized by GO terms including gliogenesis and nervous system development, and high SOX10 and SOX2 expression. Differences found between both cultures included mouse-biased Wnt signaling and ion transport genes (e.g., FXYD1) and human-biased GPCR signaling and tryptophan/indole metabolism, potentially reflecting species differences in progenitor regulation, sensory receptor repertoires, and tryptophan metabolic pathways.^88–90^ Additional differences, including HMGCS2 and RN7SL1, may relate to species-specific glial ketogenic metabolism and stress/immune responses to culturing, respectively,^84,91^ though not all differences could be functionally attributed to DRG- and species-specific biology and may also reflect differences in culture conditions.

In our DRG cultures from adult mice, coexpression of Neurog1 and Neurog2 was sufficient to induce neurons and GFAP-positive glial cells. For Neurog2, this was to be expected,^66^ but not for Neurog1, which was shown to promote differentiation of neural stem cells into solely neuronal fates.^67,92,93^ Induced neurons also expressed Brn3a, which is in line with previous studies, where Brn3a follows neurogenin (Neurog1, Neurog2) expression.^56^ Notably, co-expression of Brn3a with both neurogenins abolished the induction of GFAP-positive glial cells, likely reflecting Brn3a-mediated repression of specific non-neuronal genes,^94^ but did not increase neuronal yield compared with expression of the neurogenins alone.

We also tested Runx1 and Prdm12 for reprogramming, but these experiments were not successful. For unknown reasons, expression of both transcription factors did not result in protein abundance. It cannot be completely excluded that technical problems are the cause, but the factors themselves were readily expressed in HEK293 cells and the viral preparations were functional. Therefore, we suspect that rather the glial progenitor-like cells themselves suppressed the expression of these transcription factors.

Neurog1 and Neurog2 expression and subsequent stimulation with the neurotrophin NGF was sufficient for induction of nociceptor-like neurons in some experiments. Two weeks after infection, neurons showed immunoreactivity to nociceptor markers such as Na_V_1.9 and TrkA, as well as expression and functionality of TrpV1 and TrpA1. Overall, our data suggest that 10-30% of cells infected with Ngn2-P2A-Ngn1 have entered a nociceptor-like developmental fate, even if their electrophysiological properties did not fully match that of naïve sensory neurons. The induced nociceptive neurons were smaller in size and differed in certain electrophysiological properties when compared with naïve adult small-diameter DRG neurons in culture. Instead of multiple action potentials, induced neurons mostly generated just single spikes characterized by a shorter duration and smaller peak amplitude. Nevertheless, these transdifferentiated neurons fire action potentials and the majority of them was able to activate biphasic TTXr Na_V_ inward currents. TTXr Na_V_-mediated currents are a characteristic and important feature of nociceptors.^68,71,95,96^ This raises the question of what drives glial progenitor-like cells toward a neuronal cluster with nociceptor-like phenotypes.

A critical limitation of this study is that we were unable to reproduce the generation of nociceptor-like cells in later experiments conducted after a laboratory relocation and changes in culture conditions, including medium, serum batches, despite using the same animal facility and mouse strains. One possibility is that Neurog1 and Neurog2 act in concert with an as-yet-unidentified factor present in the culture medium to promote nociceptor differentiation. More broadly, the *in vitro* generation of sensory neurons and nociceptors remains highly variable across protocols and even between cultures within the same laboratory.^41^ It is also plausible that culture conditions alone are insufficient to drive full maturation of sensory neurons. Consistent with this, extended culturing fails to fully mature iPSC-derived sensory neurons,^97^ and adult DRG neurons in culture lose key sensory neuron hallmarks within days.^91^ Anyhow, despite extensive efforts over several years, we were unable to identify the specific variable(s) responsible for this loss of reproducibility. We therefore present these findings transparently as an initial, proof-of-principle observation that requires independent validation.

Another limitation is that we were not able to follow the fate of individual SGCs. Specific labeling of these cells, e.g. in reporter mice, is difficult because of the lack of cell type specific markers for these cells. A known marker for labeling adult SGCs is the protein Fabp7. However, SGC subtypes show variable Fabp7 expression,^98^ and our cells lost this marker when they started to divide rapidly. A large proportion of Neurog1/g2 infected cells express Sox2/Sox10, but enrichment for this failed, due to the lack of suitable biomarkers. Nevertheless, in scRNA-seq experiments, Sox2/Sox10-positive cells clustered together with immature neurons and the pseudotime analysis indicated that neurogenins induced neurogenesis from a glial progenitor cell by regulating neuronal genes over *Rest* and *Sp1*. This makes it unlikely that fibroblasts or macrophages are the target for Neurog1/Neurog2 reprogramming. The intermediate-pseudotime peak of Pax3, an early neural crest and Schwann cell precursor marker,^99^ further suggests that reprogrammed cells transiently pass through an early neural crest-like state before progressing toward a neuronal identity.

In the CNS, reactive astrocytes undergo dedifferentiation *in vivo* following gliosis.^23^ In the DRG, however, we found no evidence of gliosis as an indirect indicator of fast-dividing glial cells in the DRG after injury, neither in rodents after spared nerve injury^26^ or chronic constriction injury,^3^ nor in humans after brachial plexus injury.^13^ A gliosis-like response in the DRG has, however, been described in the context of Nageotte nodules, which form following neuronal loss, for instance in patients with diabetic peripheral neuropathy.^27^ In human DRGs from patients with neuropathic pain following brachial plexus injury, we previously observed loss of the entire neural DRG tissue in 50% of subjects.^13^ ⁸ This raises the question of whether glial progenitor-like cells with regenerative potential persist in these patients and could potentially be harnessed to promote regeneration of the adult human DRG *in vivo* after injury. Access to human DRG tissue for studies of cellular plasticity remains challenging. In the setting of brachial plexus injury, patients undergoing nerve reconstruction surgery have provided DRG tissue for research following *in vivo* functional assessment.^13^ Organ donors represent another potential source of human DRG tissue for investigating the regenerative potential and cellular plasticity of adult DRG cells (Figure 9).^83^

In summary, our findings identify adult DRG SGC-like cells as a promising endogenous source for neural reprogramming in the PNS. Their intrinsic plasticity and capacity to acquire neuronal and glial fates highlight SGCs as a potential reservoir for regenerating or repairing sensory circuits after peripheral nerve injury.

## Resource availability

### Lead contact

Requests for further information and resources should be directed to and will be fulfilled by the lead contact, Robert Blum.

### Materials availability

Plasmids generated in this study will be made available on request.

### Data and code availability

The RNA-seq data are available at GEO under accession number GSE261550 for the bulkRNA-seq and GSE261529 for the scRNA-seq data. Custom python scripts for analysis of calcium imaging and patch-clamp data are available under: https://github.com/AmSchulte/Nociceptor.

## Acknowledgements

We thank Hildegard Troll for retrovirus production. We also greatly appreciate the support from Theodore Price’s lab, particularly Andi Wangzhou, for providing us with the human DRG culture bulk-RNAseq dataset from the PRECISION Human Pain Consortium and offering valuable suggestions for its analysis.

This work was supported by the Evangelisches Studienwerk Villigst to A.S., the Interdisciplinary Center for Clinical Research Würzburg (IZKF) N-D-368 to RB and HR, and by the Deutsche Forschungsgemeinschaft (DFG) project ID: 426503586, KFO5001 ResolvePAIN to AS, HLR, RB and LE 2338/3-2 to EL. The Interdisciplinary Center for Clinical Research Würzburg (IZKF) supported the RNA-seq and corresponding data analysis in project Z-6 (FI, TB, TG) and the FACS Core Unit in project Z-12 for cell sorting. The scRNA-seq work was supported by the Bavarian Ministry of Economic Affairs, Regional Development and Energy within the project “Single cell analysis in personalized medicine“ at the Helmholtz-Institute for RNA-based Infection Research.

## Author contributions

Conceptualization: AS, HLR, RB

Methodology: AS

Investigation: AS, NK, NE, NS, AA, MK, TB, FI, RB

Software: AS

Formal analysis: AS, TB, SEJ, FI

Visualization: AS, SEJ Resources: RB, TG

Data curation: AS, TB

Funding acquisition: AS, HR, RB

Supervision: EL, HLR, RB

Project administration: AS, RB

Writing – original draft: AS, RB

Writing – review & editing: AS, HLR, RB

## Declaration of interests

The authors declare no competing or financial interests.

## STAR★Methods

### Experimental model and study participant details Cell culture and reprogramming

#### Primary DRG cell culture

Primary DRG cells were prepared from adult 4 to 6 weeks old wild-type mice as described before.^71,100^ Mice were sacrificed by CO_2_ asphyxiation and cervical dislocation. DRG were collected in ice-cold HBSS. Isolated DRG were treated with 0.6 U/ml Liberase TH (Sigma-Aldrich, Cat#5401135001, stored at 10 U/ml in dH_2_O) and 0.6 U/ml Liberase TM (Sigma-Aldrich, Cat#5401119001, stored at 10 U/ml in dH_2_O) in EDTA (3. 3̅ mM) buffered DMEM. DRG were plated on poly-L-lysine (PLL)-coated 3.5 cm culture dishes.

#### Reprogramming

Cells were cultivated in DRG medium containing DMEM/F12, GlutaMAX™ supplement (Gibco, Cat#31331028), 10% fetal calf serum (FCS, last one used: ThermoFisher Scientific, Cat# A5669501), and 1% penicillin/streptomycin supplemented with 10 ng/ml EGF (Epidermal growth factor human, Sigma-Aldrich, Cat#E9644) and FGF (Fibroblast growth factor basic human, Sigma-Aldrich, Cat#SRP6159) at 37°C, 5% CO_2_ atmosphere. After 7 d in culture, RNA was isolated or cells were split, transduced with *moloney murine leukemia virus* (MMLV)-based retroviral vectors, and plated on poly-L-lysine-coated 10 mm glass coverslips (Marienfeld, Cat#0111500). For evaluation of reprogramming efficiency, cells were fixed 7 days post infection (dpi) and used for immunocytochemistry. To differentiate the infected cells, the medium was changed from 1 to 3 dpi in 50% increments to a serum-free differentiation medium containing DMEM/F12, GlutaMAX™ supplement (Gibco), B-27 (Gibco, Cat#17504044) and N-2 supplement (Gibco, Cat#17502048), and 100 ng/ml nerve growth factor (NGF, Sigma-Aldrich, Cat#N0513). Addition of 20 ng/ml NGF, 20 ng/ml BDNF (Brain-derived neurotrophic factor, Peprotech, Cat#450-02), and 20 ng/ml GNDF (Glial cell line-derived neurotrophic factor, Peprotech, Cat#450-10) were also tested. The differentiation medium was changed weekly.

#### HEK293 cell culture

Human embryonic kidney-293 (HEK293, # ACC 305) cells were cultivated in DMEM, high glucose, GlutaMAX™ Supplement (Gibco, Cat#61965059), 10% FCS, and 1% penicillin/streptomycin at 37°C in 5% CO_2_. For testing of DNA plasmids, HEK293 cells were transfected using Lipofectamine 2000 (Thermo Fisher Scientific, Cat#11668019). Media was replaced after 24 h and cells were fixed 48 h after transfection for immunofluorescence analysis.

### Method details

#### Vector design and retroviral production

To express transcription factors, self-inactivating retroviral vectors containing a chicken beta actin promoter (pSIN-CAG backbone^101^) were cloned. Codon-optimized cDNA encoding corresponding transcription factors were produced by gene synthesis (Eurofins, ATG:biosynthetics GmbH). Transcription factor sequences were synthesized based on the protein Reference Sequence (RefSeq) of Mus musculus: neurogenin-1 (Neurog1, RefSeq: NP_035026.1); neurogenin-2 (Neurog2, RefSeq: NP_033848.1); POU domain, class 4, transcription factor 1 (Brn3a, RefSeq: NP_035273.3); runt-related transcription factor 1 isoform 1 (Runx1, RefSeq: NP_001104491.1); PR domain zinc finger protein 12 (Prdm12, RefSeq: NP_001116834.1). For bi- or tricistronic expression, inserts were linked with 2A fusion peptides (P2A: ATNFSLLKQAGDVEENPGP; T2A: EGRGSLLTCGDVEENPGP). For affinity detection, Flag (DYKDDDD), HA (YPYDVPDYA), and Myc (EQKLISEEDL) peptides were tagged to individual inserts. DNA cassettes were fused with GSG- and GGSGG-linker sequences. Red fluorescent protein (DsRed) or green fluorescent protein (GFP) served as transduction control and were expressed with the help of an IRES sequence. Constructs were cloned into CAG-DsRed^19^ and CAG-GFP^101^ backbones; Neurog2 was already available in CAG-Ngn2-DsRed.^50^ Detailed retroviral vector constructs with sources of backbone, insert, and used restriction enzymes are listed in Table S2. For validation, DNA inserts were sequenced by LGC Genomics (Berlin, Germany). Primers are listed in Table S2.

Viral vectors were produced in HEK293TN cells (SBI Biosciences) by co-transfection of the pCAG-expression plasmids and two helper plasmids with CMV-VSVG, and MMLV-CMV-gag-pol. Viral particles were purified by ultracentrifugation before application.^50^

#### RNA isolation and sequencing

For RNA isolation of 7 d old DRG cell cultures, the RNeasy Mini Kit (Qiagen, Cat#74104) was used. Cells were washed once with PBS and lysed with the provided RLT buffer supplemented with 1% β-mercaptoethanol. The cell lysate was homogenized with a 0.7 x 30 mm syringe. Cell lysates of two culture dishes were pooled. RNA was eluted in 50 µl RNAse free water.

RNA quality was checked using a 2100 Bioanalyzer with the RNA 6000 Nano kit (Agilent Technologies). The RIN for all samples was ≥8.7. DNA libraries suitable for sequencing were prepared from 500 ng of total RNA with oligo-dT capture beads for poly-A-mRNA enrichment using the TruSeq Stranded mRNA Library Preparation Kit (Illumina) according to manufacturer’s instructions (optimized for ½ volume reactions). After 15 cycles of PCR amplification, the size distribution of the barcoded DNA libraries was estimated ∼325 bp by electrophoresis on Agilent DNA 1000 Bioanalyzer microfluidic chips.

Sequencing of pooled libraries, was performed at ∼25 million reads/sample in single-end mode with 75 nt read length on the NextSeq 500 platform (Illumina). Demultiplexed FASTQ files were generated with bcl2fastq2 v2.20.0.422 (Illumina).

To assure high sequence quality, Illumina reads were quality- and adapter-trimmed via Cutadapt^102^ version 2.5 using a cutoff Phred score of 20 in NextSeq mode, and reads without any remaining bases were discarded (parameters: --nextseq-trim=20 -m 1 -a AGATCGGAAGAGCACACGTCTGAACTCCAGTCAC). Processed reads were subsequently mapped to the mouse genome (GRCm38.p6) using STAR^103^ v2.7.2b with default parameters but including transcript annotations from RefSeq annotation version 108.20200622 for GRCm38.p6. This annotation was also used to generate read counts on exon level summarized for each gene via featureCounts v1.6.4 from the Subread package.^104^ Multi-mapping and multi-overlapping reads were counted strand-specific and reversely stranded with a fractional count for each alignment and overlapping feature (parameters: -s 2 -t exon -M -O --fraction). Raw read counts per gene were converted to Transcripts Per Million (TPM) values based on combined exon length. Violin plots, and heatmap were generated using Python 3 with packages pandas and seaborn.

#### FACS sorting

Prior to FACS sorting, primary DRG cells infected with CAG-Ngn2-Ngn1-DsRed were harvested at 14 dpi in an FCS-coated Falcon. After centrifugation, cells were diluted in 500 µl of DRG medium, passed through a 100 µm cell filter (pluriSelect, Cat#43-10100-40) and placed on ice. Cell sorting and collection were performed using a BD FACS Aria III Cell Sorter (BD Biosciences). Cells were sorted based on DsRed fluorescence and collected through a 100 µm nozzle into 500 µl 0.04% BSA in 1xPBS. Cells were centrifuged and diluted to a concentration of 1,000 cells/µl in 0.04% BSA in 1xPBS prior to single cell RNA sequencing.

#### Single cell RNA sequencing

Single-cell isolation and library preparation was performed via 10x Genomics Chromium Next GEM 3’ kit v3.1 following the manufacturer’s instruction. In brief, FACS-sorted cells were subsequently loaded on a 10x Genomics Chip G and encapsulated into Gel Bead-in-Emulsion (GEM) droplets, each containing a single cell along with a unique barcoded bead. Within these GEMs, the mRNA from each cell was reverse transcribed into complementary DNA (cDNA) using reverse transcriptase, with the process occurring on the uniquely barcoded beads. The resulting cDNA was then recovered from the GEM droplets and amplified to prepare sequencing libraries using PCR. During library preparation, additional sequences necessary for sequencing, such as adapter sequences, were incorporated into the cDNA fragments. Final libraries were sequenced on an Illumina NextSeq 2000 platform.

#### Indirect immunofluorescence staining

Cells were fixed with 4 % paraformaldehyde (PFA) in phosphate buffer for 15 min at 37°C and stored in 1xPBS at 4°C when not directly processed. For cell permeabilization and blocking, cells were incubated with 10% horse serum (Sigma-Aldrich, Cat#H1270), 0.1% Triton X100, and 0.1% Tween 20 in 1xPBS (blocking solution) for 1 h at RT. Primary antibodies in blocking solution were incubated for 2 h at RT. After washing the cells 8 times with 0.1% Triton X100, and 0.1% Tween 20 in 1xPBS (washing solution), secondary antibodies in a concentration of 0.5 μg/ml in blocking solution were incubated for 1.5 h at RT. Used antibodies are listed in Table S3. Cells were washed again with washing solution and 1xPBS, and nuclei were stained with 0.4 μg/ml DAPI in 1x PBS. Finally, cells were washed with 1xPBS, dipped in water, dried, and embedded in Aqua-Poly/Mount (Polysciences, Cat#18606).

#### Microscopy

Immunocytochemical stainings and tile scans were captured with an Axio Imager M2 (Zeiss) using a Plan Apochromat 20x (air, numerical aperture (NA): 0.8). For higher resolution, an inverted Olympus IX81 microscope combined with an Olympus FV1000 confocal laser scanning system was used. Images were acquired with an FVD10 SPD spectral detector (Olympus) and diode lasers of 405, 473, 559, and 635 nm using an Olympus UPLAPO 20x (air, NA: 0.75), Olympus UPLFLN40x (oil, NA: 1.3), or UPLSAPO60x (oil, NA: 1.35). The pinhole setting represented one Airy disc. 12-bit greyscale images were processed with maximum intensity projection, adjusted in brightness and contrast, and merged into an RGB composite using Fiji.^105^ Final processing was done with Adobe Photoshop CS5 (Adobe Systems, RRID:SCR_014199).

#### Calcium imaging

Imaging of cytosolic calcium was performed with Oregon Green 488 BAPTA-1 AM (OGB1, Invitrogen, Cat#O-6807). Cells were loaded with 5 µM OGB1 for 15 min at 37 °C in HEPES-buffered ACSF (artificial cerebrospinal fluid, 120 mM NaCl, 2.5 mM KCl, 1.2 mM MgCl_2_, 2.4 mM CaCl_2_, 1.2 mM NaH_2_PO_4_, 26 mM NaHCO_3_, 10 mM Glucose, 10 mM HEPES). 5 mM OGB1 (in 20% pluronic/DMSO) was stored in small single-use aliquots.

The imaging setup for fast calcium imaging consisted of a BX51WI upright microscope (Olympus) equipped with a water-immersion objective (20× Olympus UPLFLN, NA 0.5) and a pE-4000 fluorescence illumination system (CoolLED). Images were captured at 5 Hz with a Rolera XR Mono fast 1394 CCD camera (Qimaging) controlled by the streaming software Streampix 4.0 (NorPix). A camera binning of 2×2 was used.

Cells expressing Ngn2-P2A-Ngn1-^IRES^DsRed were imaged 14 and 15 dpi at room temperature in a plastic perfusion chamber in HEPES-buffered ACSF under continuous perfusion with the help of a Minipuls 3 peristaltic pump (speed of the perfusion pump: 15 A.U., purple tubing, Gilson). The DsRed signal excited by 550 nm wavelength was captured to identify infected cells. Changes in calcium signals (OGB1: 470 nm excitation) were imaged in response to 10 µM capsaicin (Sigma-Aldrich, Cat#M2028) and 100 µM allyl-isothiocyanate (AITC, Sigma-Aldrich, Cat#36682). Capsaicin was applied for 10s through perfusion with the peristaltic pump, and 1 ml AITC was applied for 10s with a pipette.

#### Electrophysiology

For electrophysiological measurements of reprogrammed neurons, current-clamp and voltage-clamp recordings were performed in the whole-cell configuration at room temperature. Extracellular solution for current-clamp and paired pulse voltage-clamp recordings contained (in mM): 120 NaCl, 3 KCl, 2.5 CaCl_2_, 1 MgCl_2_, 30 HEPES, 15 glucose (pH 7.4 with NaOH); the corresponding intracellular solution contained 125 KCl, 8 NaCl, 1 CaCl_2_, 1 MgCl_2_, 0.4 Na_2_-GTP, 4 Mg-ATP, 10 EGTA, 10 HEPES (pH 7.3 with KOH).^73^ The extracellular solution used to characterize specifically TTX-resistant, Na_V_-mediated currents contained (in mM): 150 NaCl, 2 KCl, 1.5 CaCl_2_, 1 MgCl_2_, 10 HEPES, 0.0005 TTX (pH 7.4 with NaOH); and the pipette contained (in mM): 35 NaCl, 105 CsF, 10 EGTA, 10 HEPES (pH 7.3 with CsOH).^73^ Reprogrammed (14-17 dpi) or naive (DIV 1-2) neurons were continuously perfused with the help of a Minipuls 3 peristaltic pump (speed of the perfusion pump: 3 A.U., purple tubing, Gilson). Pipettes with a resistance of 2−4 MΩ were pulled from borosilicate glass (Science Products, Cat#GB 150-8P). Data were acquired using a HEKA EPC-10 USB patch-clamp amplifier (HEKA Electronic, Reutlingen, Germany) controlled by the PatchMaster software (v2x90, HEKA Electronic, RRID:SCR_000034) at a sampling interval of 40 μs. Voltages were not corrected for the liquid junction potential. Data were processed with Fitmaster 2x91 (HEKA Electronic, RRID:SCR_016233) and analyzed using python scripts (https://github.com/AmSchulte/Nociceptor).

#### Current-clamp recordings

After establishing the whole-cell configuration, cells were voltage-clamped at a holding potential of -100 mV and tested for functional ion channel expression by measuring inward and outward currents with a 50 ms test pulse to -20 mV followed by a 100 ms test pulse to 60 mV, respectively. The resting membrane potential (RMP) was measured at zero current injection immediately after switching to the current-clamp mode. Subsequently, cells were clamped to -60 mV using the low-frequency voltage clamp available in the PatchMaster software. Action potentials were evoked by a series of current injections increasing from 0 to 220 pA in steps of 20 pA. The duration of current injections was set to 3 ms to evoke single action potentials or to 300 ms to trigger trains of action potentials. Data were low-pass filtered at 3 kHz. Evoked events were considered action potentials if their peak voltage exceeded 0 mV and a firing threshold could be detected for the first action potential. The firing threshold of an action potential was defined as a voltage at which d*V*/d*t* reached the level of 0.03 × (d*V*/d*t*_max_−d*V*/d*t*_min_) + d*V*/d*t*_min._ The spike width was analyzed at a fixed voltage of -20 mV.

#### Voltage-clamp recordings

Cells were held at -130 mV for at least 8 min to facilitate recovery of Na_V_ channels from inactivation. Activation of Na_V_ channels was measured with test depolarizations between -110 and 30 mV in steps of 10 mV, delivered every 10 s. The test pulse duration was 100 ms. Leak and capacitive currents were measured with a p/n method and subtracted online. Data were low-pass filtered at 3 kHz.

### Quantification and statistical analysis

#### Comparisons of bulk RNAseq with a published single cell RNAseq dataset

Published single cell datasets of DRG cells^35^ are available as a web-based portal at Broad Institute (https://singlecell.broadinstitute.org/single_cell/study/SCP1539/). The raw data is deposited at Gene Expression Omnibus: GSE174430 (Cell_acute data) and GSE188971 (Cell_culture data).

#### Comparison to pseudo bulk datasets

The bulk analysis of the cultured DRG cells was compared to the single cell datasets by transforming the single cell datasets into pseudo bulk datasets via the R package SingleR.^106^ Next, SingleR is used to compare the bulk dataset to the two pseudo bulk datasets using the gene expression in the pseudo bulk datasets as a reference. The output is a Pearson correlation with 0 indicating no correlation and 1 indicating perfect positive correlation.

#### Comparison to cell types in single cell RNAseq datasets

Markers for the cell types in the acute single cell dataset were identified with the R package Scran^107^ by comparing the gene expression in each individual cell type with all the other cell types. Heatmaps showing the expression levels in the bulk dataset of the cell type markers were made in R using the R package Scater.^108^

#### Comparison of bulk-RNAseq of mouse and human DRG cultures

As a human DRG culture bulk RNA dataset, we used a published bulk-RNAseq dataset from^83^ (GEO accession: GSE300240). The three vehicle control samples from “GSE300240_Normalized_Counts_GEO_submission.csv“ were used as the human DRG culture dataset (day 8 of culture). Human Ensembl gene identifiers were mapped to gene symbols using the GENCODE v38 annotation. Our mouse dataset comprised 6 replicates of DRG cultures at day 7 in culture, quantified as gene-level read counts.

The 1000 most highly expressed genes were extracted from each dataset (by mean normalized counts for human and mean read counts for mouse) and tested for enriched ontology terms with Metascape,^109^ analyzed as species *Mus musculus*.

Within each dataset, gene-level expression was log-transformed (log1p) and z-standardized per sample (across all genes) to zero mean and unit standard deviation. For cross-species comparison, high-confidence one-to-one orthologs were retrieved from Ensembl BioMart (Ensembl release 116, GRCh38.p14/GRCm39).^110^ Only pairs annotated as ortholog_one2one with an orthology confidence of 1 and non-missing identifiers on both sides were retained, yielding 16,032 ortholog pairs. After matching the gene names to both datasets, 15,476 genes remained. The global transcriptional concordance between the human and mouse cultures was assessed by Spearman’s rank correlation of per-gene mean expression. To identify species-biased genes, we computed the human–mouse residual (mouse z − human z) and selected the 300 genes with the strongest bias in each direction. Enriched GO Biological Process terms for each set were identified with EnrichR.^111^ Unless stated otherwise, all analyses were performed in Python 3.13.2. Figures were generated with matplotlib (v3.10.1) and seaborn (v0.13.2).

#### Single cell RNA sequencing analysis

Single cell RNAseq data analysis was performed using the scanpy package (version 1.9.2) in python 3. For quality control, genes that are detected in less than 3 cells and cell with less than 200 genes were filtered out. In addition, cells with more than 20% of mitochondrial transcripts and more transcripts than the 98% quantile in total were removed. After normalization to 10,000 reads per cell and logarithmization, highly variable genes were extracted using default parameters. Effects of total counts per cell and the percentage of mitochondrial genes expressed were regressed out, the data were scaled to unit variance, and values exceeding standard deviation of 10 were clipped. To reduce the dimensionality of the data, principal component analysis (PCA) was performed on the most variable genes. Using the PCA representation of the data matrix, a neighborhood graph was computed using 15 neighbours and 15 principal components. Uniform manifold approximation and projection (UMAP) was used to embed and visualize the graph in two dimensions. Leiden clustering was performed with a resolution of 0.17 and marker genes were determined using the Wilcoxon method.

The “Unidentified” cluster was excluded from further analysis. The official Python 3 interface (gprofiler-official 1.0.0) to the g:Profiler toolkit was used for enrichment analysis of functional GO terms corresponding to *Mus musculus* based on the top 200 differentially expressed marker genes per cell cluster. Cluster were compared to the developmental stages of somatosensory neurons analyzed in a scRNA-seq study by Sharma et al.^61^, ranging from E11.5 to adulthood. Top 50 enchriched genes of the six developmental time points were used for running the univariate linear model (ULM) implemented in decoupler (version 2.2.0),^112^ and the mean ULM score per cell cluster was calculated. For trajectory analysis, the neighborhood graph was recomputed, and a diffusion map was calculated. This was followed by neighborhood graph construction from the diffusion map embedding, and partition-based graph abstraction (PAGA)^113^ using scanpy. To analyze the transition from “Glial progenitors” to “Immature neurons”, the trajectory analysis was recomputed using only these two clusters and a pseudotime and transcription factor activity analysis was performed. The diffusion pseudotime (DPT)^114^ was calculated using the *Glial progenitors* cluster as the root population. Transcription factor activities were inferred from the mouse CollecTRI regulatory network^115^ by running ULM in decoupler. Transcription factors were ranked according to the association of their inferred activity with pseudotime using distance correlation, and the highest-ranking factors were selected for downstream visualization. Expression dynamics of transcription factor target genes along pseudotime were visualized using decoupler.

#### Image analysis

Whole cell culture coverslips stained for neuronal and glial markers were tile-scanned with 14 bit at a resolution of 0.454 µm/ pixel. Analysis of the large amount of data (5-6 GB per cover) was performed with CellProfiler.^116^ Each cover scan was processed in a batch of tif-converted single-tile images.

Quantification of DAPI, Tuj1, Map2, GFAP, or Brn3a-positive cells was performed with the “IdentifyPrimaryObjects” module. Here, the minimum cross entropy thresholding method was used, where the lower bounds on the threshold needed to be adjusted manually for each cover scan to counteract staining variability. The RFP-positive area was defined with a manual threshold and the resulting mask was projected onto the marker staining (module: “MaskImage”) to determine how many of the infected cells were positive for the respective marker. Results were exported to spreadsheets and further processed and visualized in python.

For quantification of Sox2 and Sox10-positive infected cells, tile scans consisting of 120 single images were captured and Stardist was used to quantify Sox2, Sox10, or DAPI-stained cell nuclei. For RFP quantification, Otsu thresholding and a median filder of 3-pixel disk size was applied. Nuclei overlapping with the RFP mask were regarded as infected cells positive for the respective marker. Only cell bigger that 100 pixels were taken into account.

#### Analysis of calcium imaging signals

Calcium imaging videos of cells stimulated with either Capsaicin or AITC were analyzed in python. Region of interest (ROI) detection was performed with the open-source DL model StarDist2D.^117^ First, the DsRed signal was segmented with StarDist2D to detect infected cells. Based on the resulting ROIs, calcium imaging signal traces were extracted for each infected cell. For uninfected cells, the DsRed-positive ROIs were removed from the calcium imaging video, and ROI detection and signal extraction was performed on the rest of the OGB1-stained cells. To identify the proportion of reacting cells, signal traces were low-pass filtered with a wavelet function. When the derivative of the low-pass filtered signal reached a uniform threshold, the cell was counted as reacting. The proportion of reacting infected cells was compared to the proportion of reacting uninfected cells. The analysis script is openly available (https://github.com/AmSchulte/Nociceptor).

#### Statistical analysis

Data were tested for significance using python 3 with packages scipy.stats and scikit-posthocs. The normal distribution of the data was assessed with the Shapiro test. Non-normally distributed data were compared using the Kruskal-Wallis H-test with post hoc Mann-Whitney U test for more than two groups, or a two-sided Mann-Whitney U test when comparing two groups.

Normally distributed data were tested first for equal variance with the Bartlett’s test. More than two groups were tested with the Alexander Govern test when the variance was unequal, or with the one-way ANOVA when the variance of the data was equal. Then, a post hoc T-test was performed. Two groups with unequal variance were compared with the Welch’s t-test, or with the unpaired, two-tailed t-test when having an equal variance. Bonferroni correction was applied for multiple comparisons. Data are presented as the mean ± standard error of the mean (s.e.m.). A p-value < 0.05 was considered statistically significant. In boxplots, single data points are presented. The boxes extend from the 25^th^ to the 75^th^ percentile and the lines within the boxes show the median. The whiskers extend from the smallest to the highest value, except for outliers.

## Key resources table

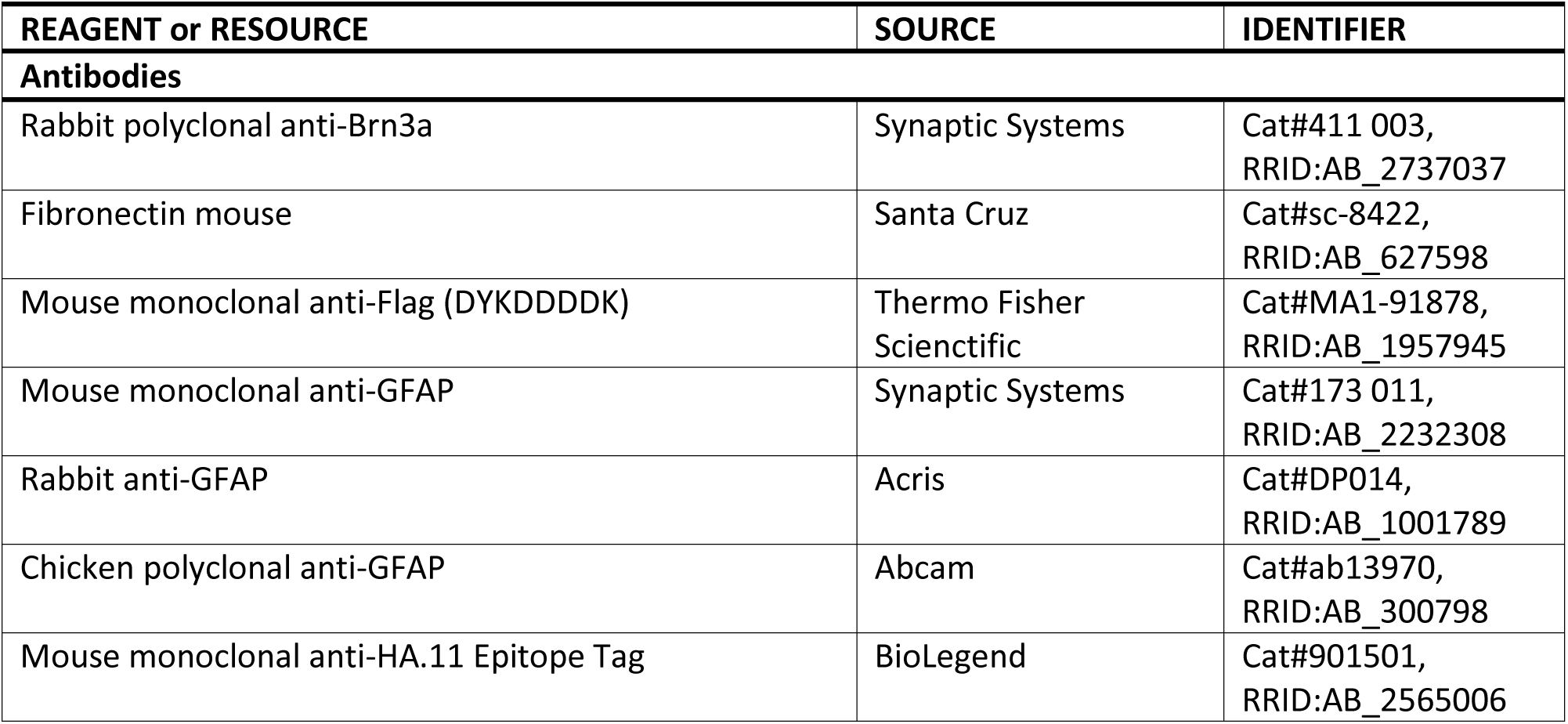
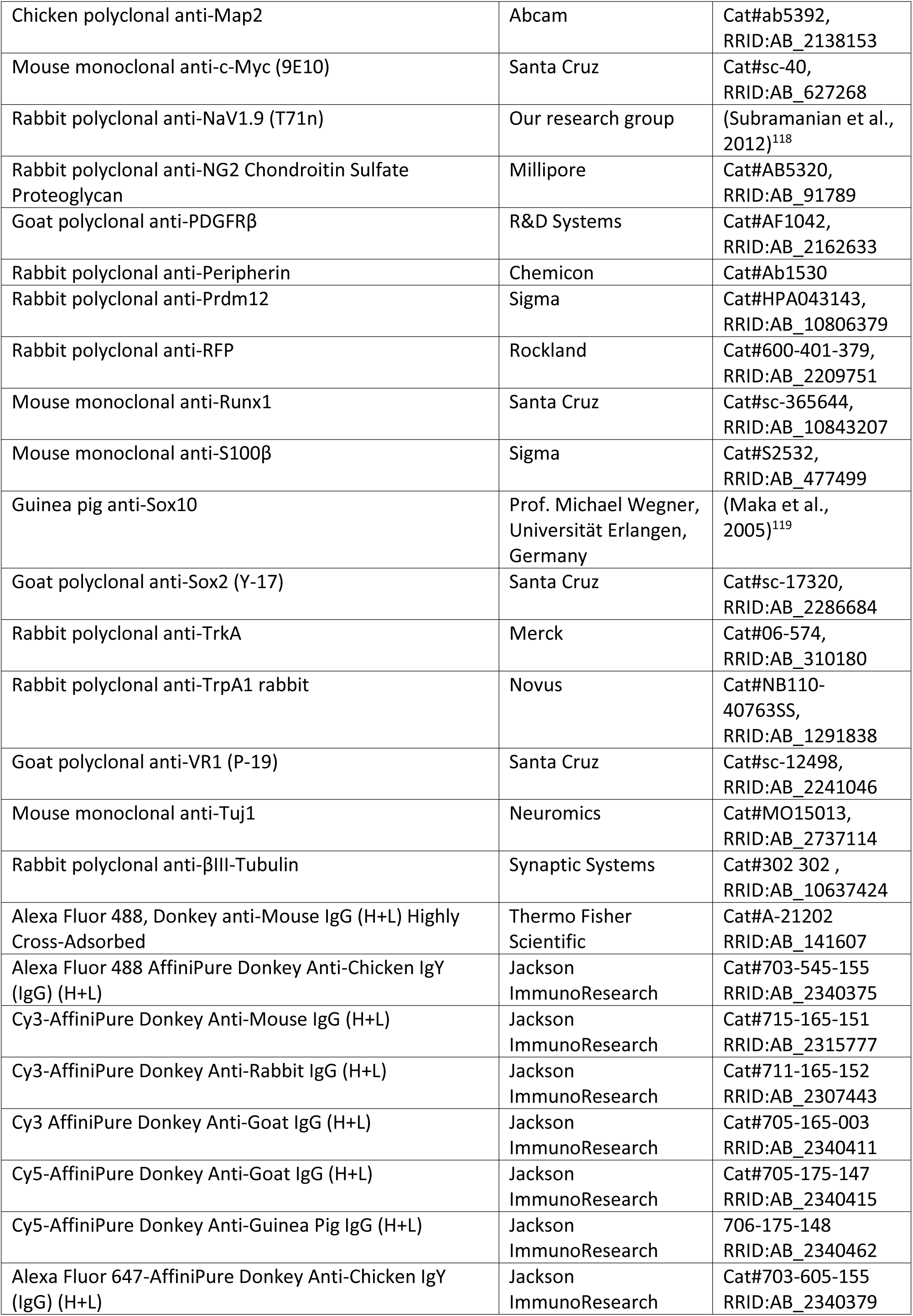
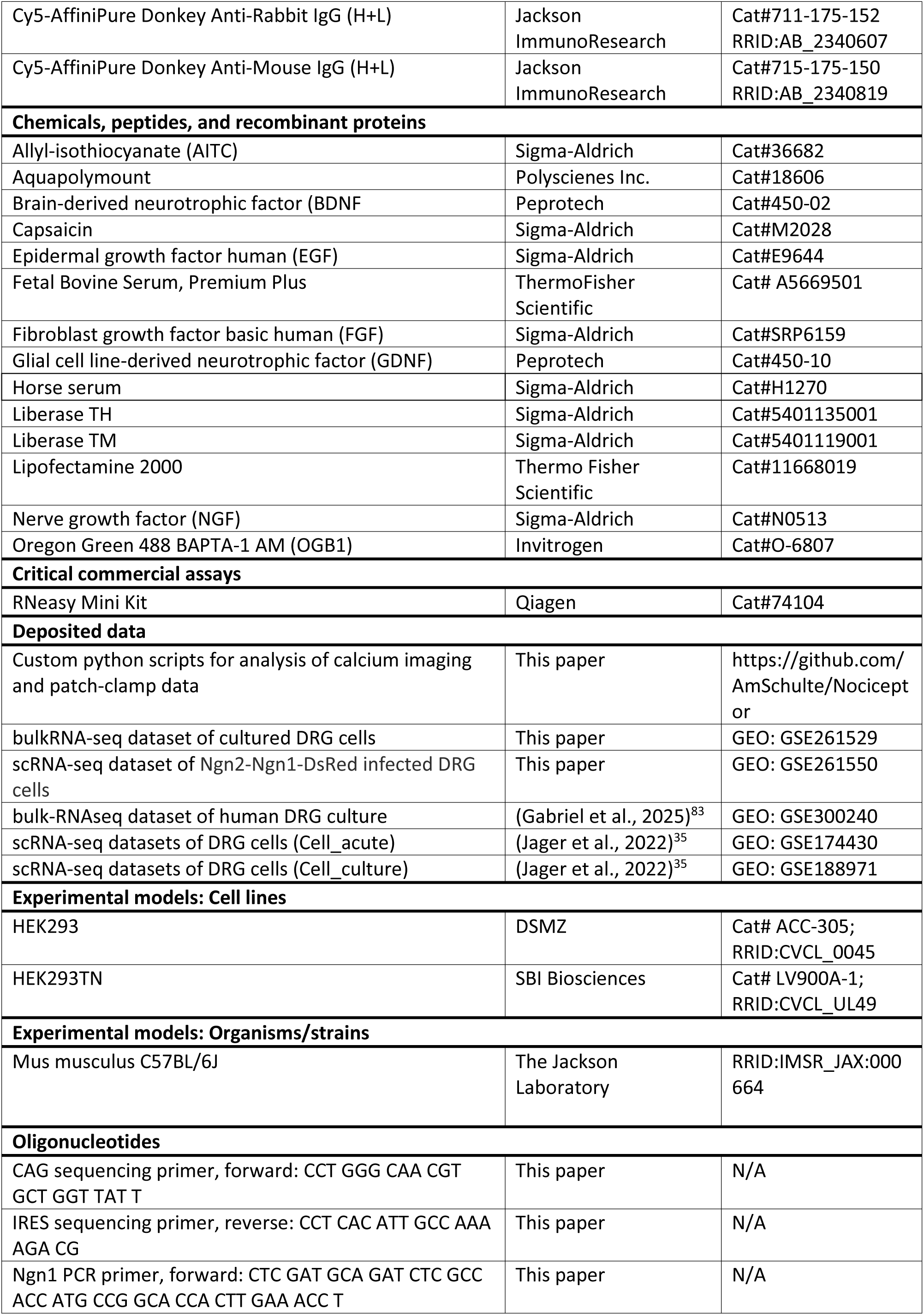
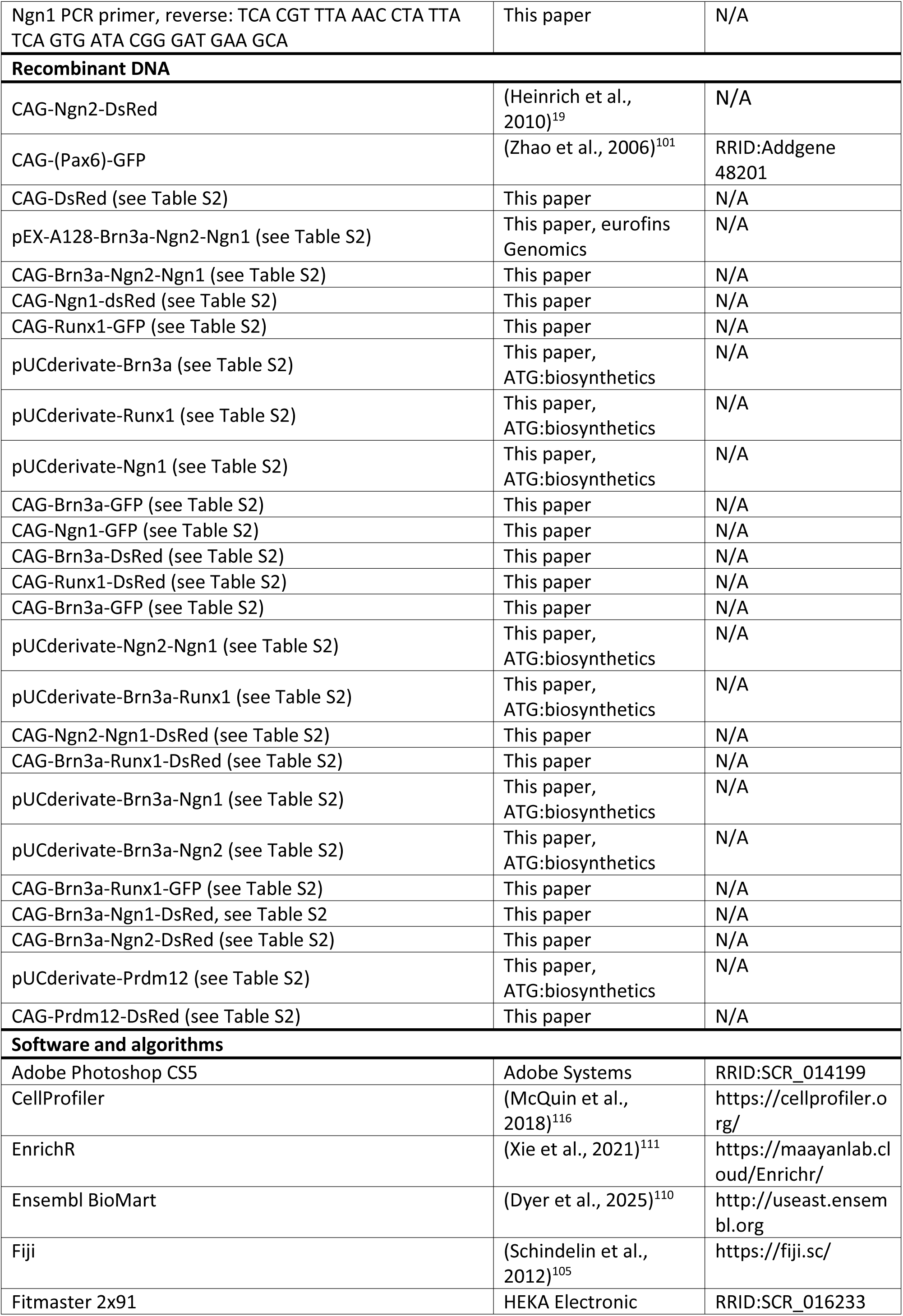
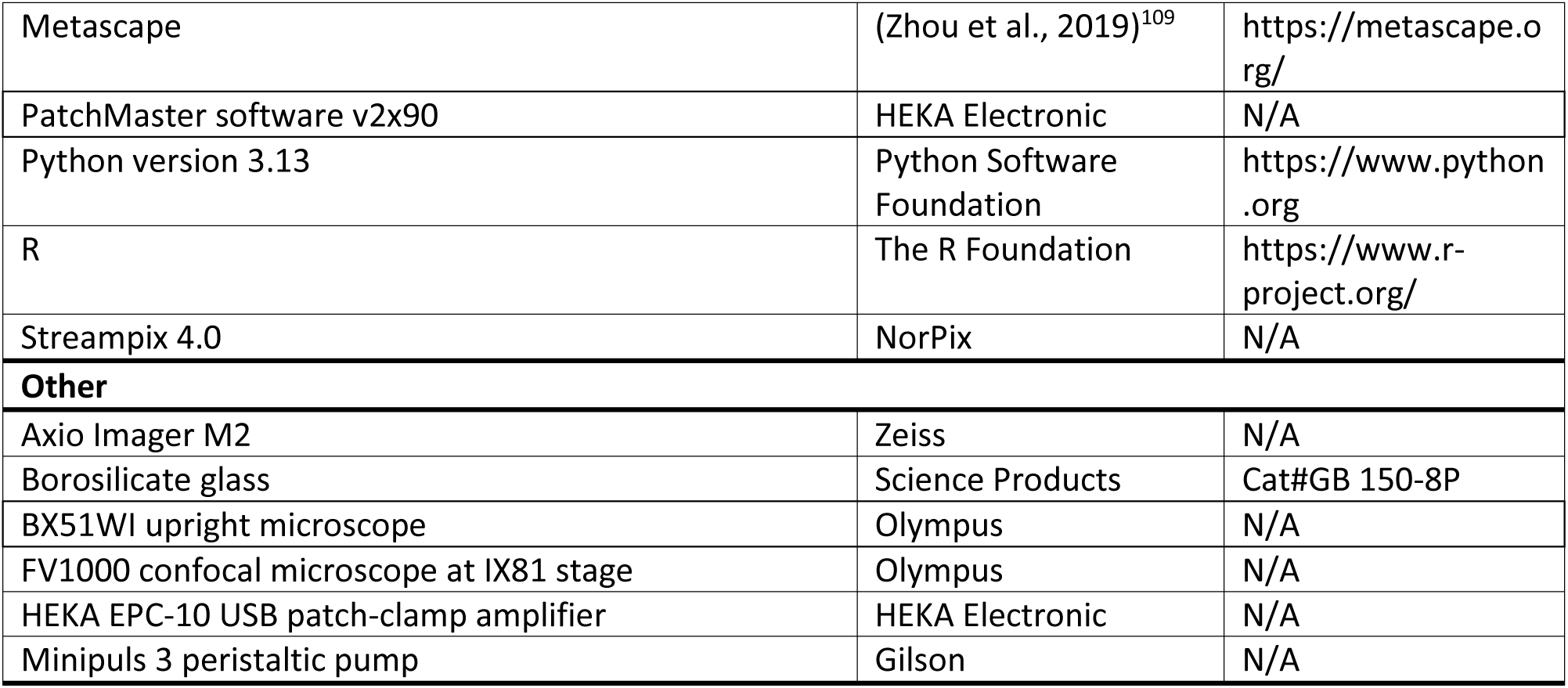

## Supplemental information

Fig. S1. Sensory progenitor markers are expressed at similar levels as housekeeping genes.

Fig. S2. Retroviral vectors transfected and expressed in HEK293 cells.

Fig. S3. Retroviral vectors with no effect on peripheral glial progenitor-like cells.

Fig. S4. Variability of reprogramming outcome after ^IRES^DsRed-Ngn2-P2A-Ngn1 induction.

Table S1. TPM count table of bulk RNAseq of mouse DRG culture at DIV 7.

Table S2. Transfer plasmids for retroviral vectors based on the pSIN-CAG backbone

## Notes

### Competing Interest Statement

The authors have declared no competing interest.

### Summary of Updates

Title changed; Abstract/Summary updated; Introduction updated; added genes to Fig. 1 E ; Fig. 7 revised; Fig. 8 (previously Fig. S4) updated; Fig. 9 added; new Fig. S4 added, Table S1 (TPM table) added; Results updated; Discussion updated.

