## Supplemental Information for "Adult dorsal root ganglion glial cells exhibit neurogenic potential upon reprogramming"

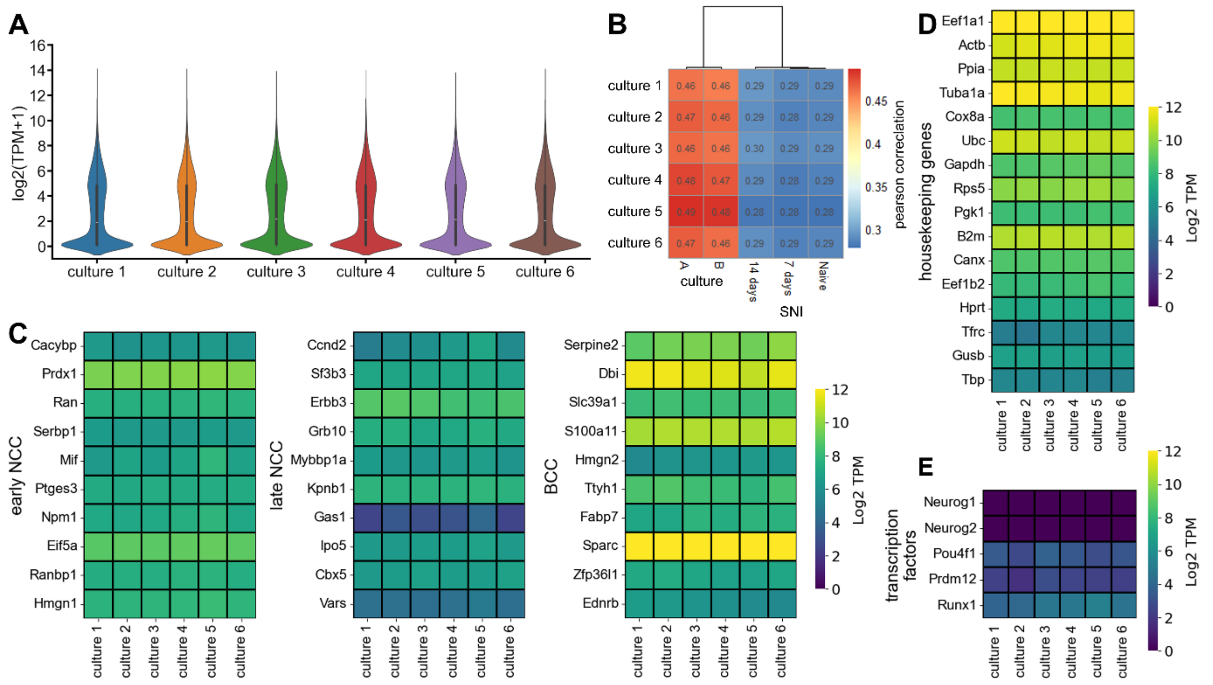


**Fig. S1. Sensory progenitor markers are expressed at similar levels as housekeeping genes**

(A) TPM count distribution for biological replicates of RNA sequenced DRG cell culture (DIV 7) shown as violin plots.

(B) Comparison of the bulk dataset to pseudo bulk datasets of cultured (A and B culture) or acutely dissociated DRG of naïve and SNI treated mice (Naïve, 7 days, 14 days) (Jager et al., 2022)28. Scale bar: Pearson correlation.

(C) Transcriptomic expression of the top 10 differentially expressed genes of early and late NCC (neural crest cells) and boundary cap cells (BCC) (Faure et al., 2020)47.

(D) Transcriptomic expression of housekeeping genes in DRG cell culture (DIV 7).

(E) Expression profile of transcription factors used for reprogramming. Each column represents a sample of a DRG cell culture (DIV 7) of adult wt mice (n = 6). Scale bar: Log2 transcripts per million (TPM).

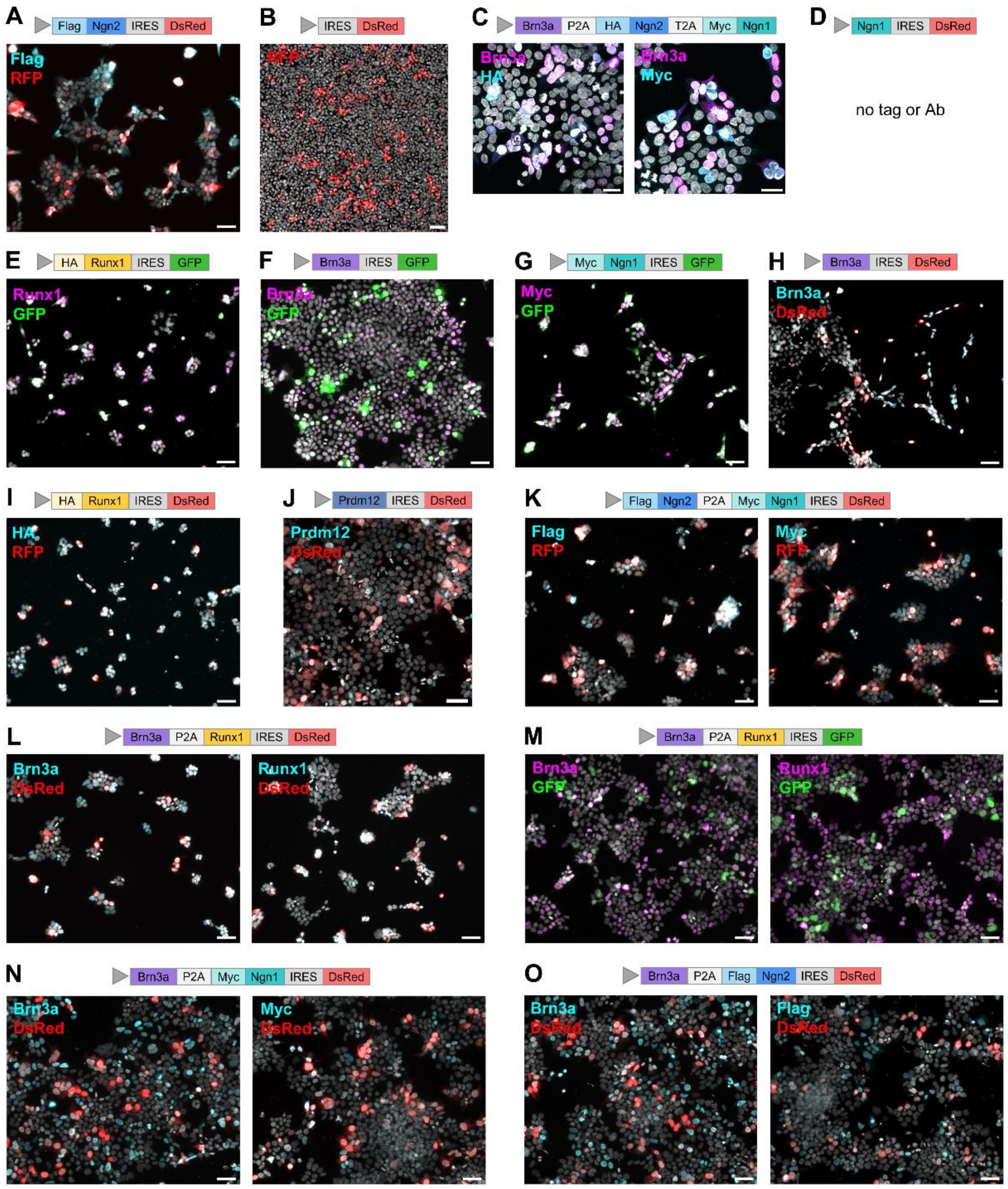


**Fig. S2. Retroviral vectors transfected and expressed in HEK293 cells**

(A) Expression of the CAG-Flag-Ngn2-IRES-DsRed vector (Heinrich et al., 2010)11.

(B) Its CAG-IRES-DsRed backbone served as a control.

(C) Expression of the triple vector with Brn3a (violet), Neurog2 (Ngn2, HA-tagged, blue), and Neurog1 (Ngn1, Myc-tagged, cyan) sequences.

(D-I) Neurog1, Runx1 (HA-tagged, yellow), and Brn3a were expressed with IRESDsRed or IRESGFP vectors. (J) CAG-Prdm12-IRES-DsRed enabled Prdm12 (grey-blue) expression in HEK293 cells.

(K-O) HEK293 cells expressing the bicistronic constructs Flag-Ngn2-P2A-Myc-Ngn1 (K), Brn3a-P2A-Runx1 (L, M), Brn3a-P2A-Myc-Ngn1 (N), or Brn3a-P2A-Flag-Ngn2 (O).

HEK293 cells expressing retroviral vectors were labeled 48-72h after transfection. The grey arrowhead represents the CAG promotor. Except for the triple vector, mono- and bicistronic constructs contained DsRed or GFP as a fluorescent protein marker. Scale bars: 50 µm (C: 25µm). GFP, green fluorescent protein.

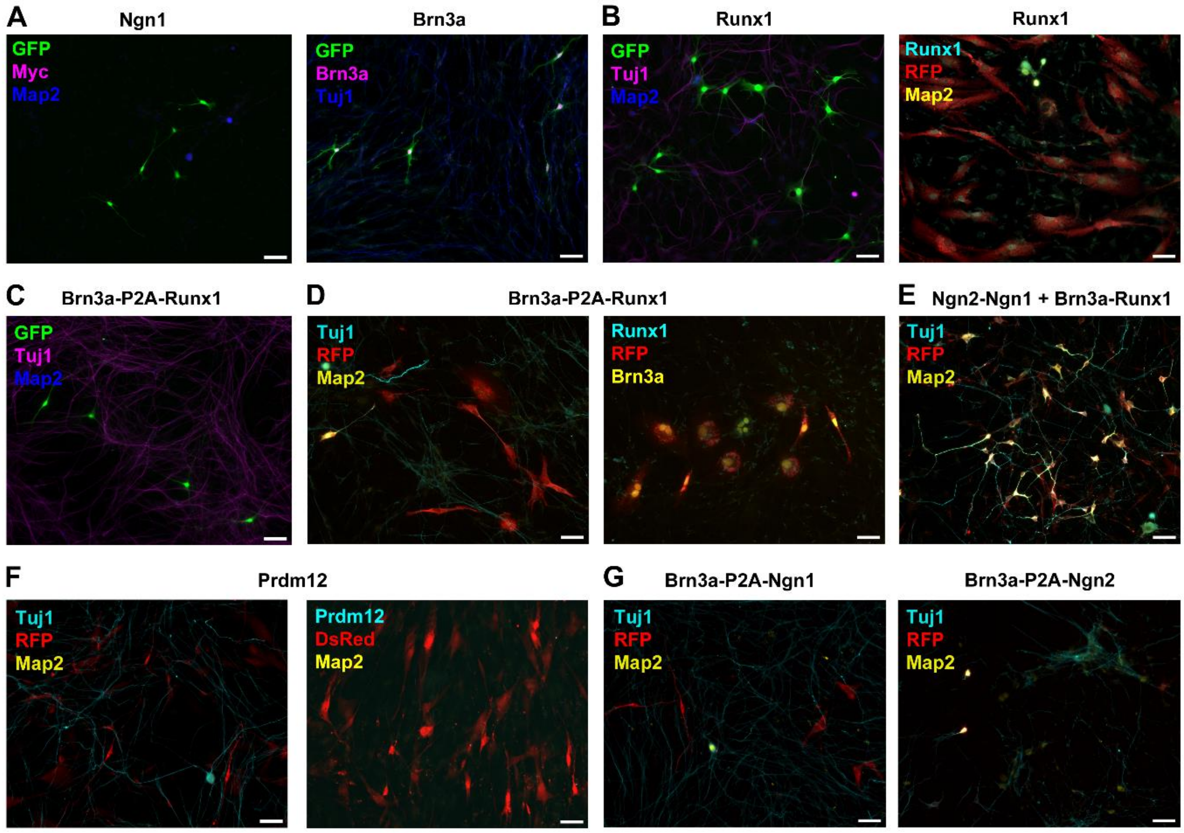


**Fig. S3. Retroviral vectors with no effect on peripheral glial progenitor-like cells**

Representative images of glial progenitor-like cells transduced with indicated retroviral vectors at 7 dpi. IRESGFP (A-C) or IRESDsRed (stained: RFP) (B, D-G) served as infection markers. The neuronal markers βIII-Tubulin (Tuj1) and Map2 were used to visualize if a neuronal phenotype was induced. Scale bars: 50 µm.

(A) Expression of Neurog1 (Ngn1, Myc-tagged, magenta) or Brn3a (magenta).

(B) Expression of Runx1 (cyan).

(C-D) Coexpression of Brn3a and Runx1.

(E) Coexpression of Brn3a and Runx1 in combination with the Ngn2-P2A-Ngn1 vector.

(F) Expression of Prdm12 (cyan).

(G) Coexpression of Brn3a with Neurog1 or Neurog2.

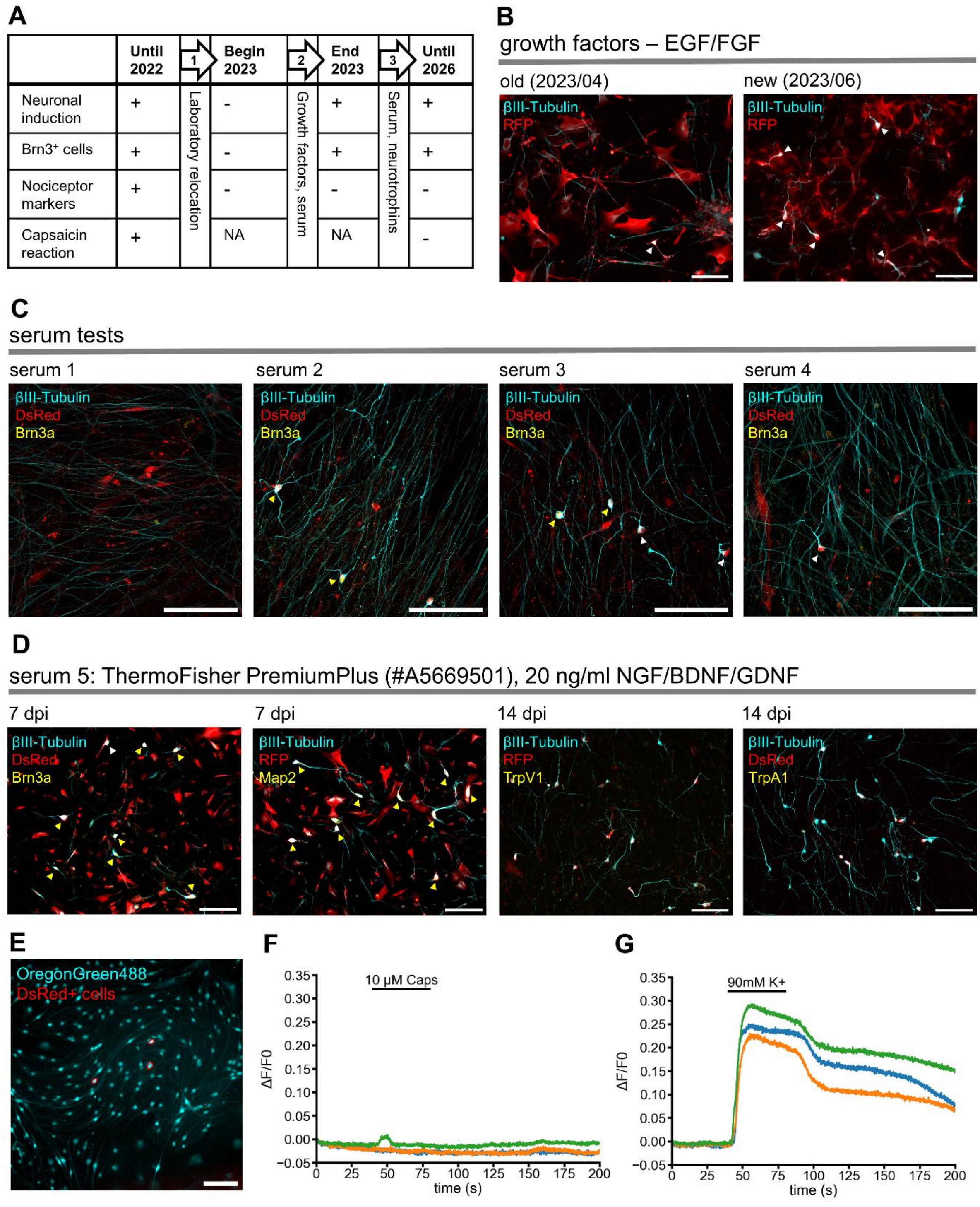


**Fig. S4. Variability of reprogramming outcome after IRESDsRed-Ngn2-P2A-Ngn1 induction**

(A) Characteristics of reprogrammed cells over several years changed depending on (un)known factors. Arrows mark three time points where certain culture parameters changed.

(B) Representative immunofluorescence staining of reprogrammed cells before and after addition of a new batch of the growth factors EGF and FGF. Cells are labelled against βIII-Tubulin (cyan) for neurons and RFP (red) for IRESDsRed-positive infected cells at 7 days post infection (dpi). Induced neurons are marked with white arrowheads.

(C) Test of different types of serum evaluated by induction of βIII-Tubulin-positive neurons (white arrowhead) and Brn3a-positive neurons (yellow arrowhead).

(D) Reprogrammed cells after addition of Thermofisher PremiumPlus serum and the neurotrophic factors NGF, BDNF, and GDNF labelled for βIII-Tubulin-positive neurons (cyan) and DsRed-positive infected cells (red). Induced Brna3- or Map2-positive neurons are marked with yellow arrowheads at 7dpi. At 14 dpi, cells were labelled for TrpV1 or TrpA1 (yellow).

(E) Average calcium signals detected with Oregon Green 488 (cyan) in IRESDsRed-Ngn2-P2A-Ngn1 infected cells (15 dpi), encircled in red.

(F-G) Calcium signals of three IRESDsRed-positive cells stimulated with 10 µM Capsaicin (F) or 90mM K+ (G) for 40s.

Scale bars: 100 µm.

**Table S2. Transfer plasmids for retroviral vectors based on the pSIN-CAG backbone.**

| Vector (#internal ID) | Detailed construct | Source | |
|---|---|---|---|
| | | **Backbone; Enzyme** | **Insert; Enzyme** |
| CAG-Ngn2-DsRed (#576) | CAG-Kozak-ATG-Flag-Ngn2-IRES-DsRed | Heinrich et al., 2010 | |
| CAG-(Pax6)-GFP (#582) | CAG-Kozak-ATG-(Pax6)-IRES-GFP | Zhao et al, 2006; RRID:Addgene 48201 | |
| CAG-DsRed (#955) | CAG-IRES-DsRed | CAG-Ngn2-DsRed (#576); BamHI, PmeI | none |
| pEX-A128-Brn3a-Ngn2-Ngn1 (#976) | Kozak-ATG-Brn3a*-GSG-P2A-GSG-HA-GGSGG-Ngn2*-G^^SG-T2A-GSG-Myc-GGSGG-Ngn1* | gene synthesis, eurofins Genomics | |
| CAG-Brn3a-Ngn2-Ngn1 (#1013) | CAG-Kozak-ATG-Brn3a*-GSG-P2A-GSG-HA-GGSGG-Ngn2*-GSG-T2A-GSG-Myc-GGSGG-Ngn1* | CAG-Ngn2-DsRed (#576); BamHI, PmeI (blunt) | pEX-A128-Brn3a-Ngn2-Ngn1(#976); EcoRV (blunt) |
| CAG-Ngn1-dsRed (#1025) | CAG-Kozak-ATG-Ngn1*-IRES-DsRed | CAG-Ngn2-DsRed (#576); BamHI, PmeI | Ngn1 PCR ampliflied from pEX-A128-Brn3a-Ngn2-Ngn1(#976); BglII, PmeI |
| CAG-Runx1-GFP (#1038) | CAG-Kozak-ATG-HA-GSG-Runx1*-IRES-GFP | CAG-Pax6-GFP (#582); BamHI | pUCderivate-Runx1 (#1041); BglII |
| pUCderivate-Brn3a (#1040) | Kozak-ATG-Brna3* | gene synthesis, ATG:biosynthetics | |
| pUCderivate-Runx1 (#1041) | Kozak-ATG-HA-GSG-Runx1* | gene synthesis, ATG:biosynthetics | |
| pUCderivate-Ngn1 (#1042) | Kozak-ATG-Myc-GSG-Ngn1* | gene synthesis, ATG:biosynthetics | |
| CAG-Brn3a-GFP (#1043) | CAG-Kozak-ATG-Brn3a*-IRES-GFP | CAG-Pax6-GFP (#582); BamHI | pUCderivate-Brn3a (#1040); BamHI |
| CAG-Ngn1-GFP (#1044) | CAG-Kozak-ATG-Myc-GSG-Ngn1-*IRES-GFP | CAG-Pax6-GFP (#582); BamHI | pUCderivate-Ngn1 (#1042); BamHI |
| CAG-Brn3a-DsRed (#1045) | CAG-Kozak-ATG-Brn3a*-IRES-DsRed | CAG-Ngn2-DsRed (#576); BamHI, PmeI (blunt) | pUCderivate-Brn3a (#1040); BamHI (blunt) |
| CAG-Runx1-DsRed (#1051) | CAG-Kozak-ATG-HA-GSG-Runx1*-IRES-DsRed | CAG-Ngn2-DsRed (#576); BamHI, PmeI (blunt) | pUCderivate-Runx1 (#1041); BglII (blunt) |
| CAG-Brn3a-GFP (#1056) | CAG-Kozak-ATG-Brn3a*-IRES-GFP | CAG-Brn3a-DsRed (#1045); HindIII, NotI | CAG-Brn3a-GFP (#1043); HindIII, NotI (GFP exchanged) |
| pUCderivate-Ngn2-Ngn1 (#1057) | Kozak-ATG-Flag-GSG-Ngn2*-GSG-P2A-GSG-Myc-GGSGG-Ngn1* | gene synthesis, ATG:biosynthetics | |
| pUCderivate-Brn3a-Runx1 (#1058) | Kozak-ATG-Brn3a*-GSG-P2A-GGSGG-Runx1* | gene synthesis, ATG:biosynthetics | |

| | | | |
|---|---|---|---|
| CAG-Ngn2-Ngn1-DsRed (#1063) | CAG-Kozak-ATG-Flag-GSG-Ngn2*-GSG-P2A-GSG-Myc-GGSGG-Ngn1*-IRES-DsRed | CAG-Ngn2-DsRed (#576); BamHI, PmeI | pUCderivate-Ngn2-Ngn1 (#1057); BglII, EcoRV |
| CAG-Brn3a-Runx1-DsRed (#1064) | CAG-Kozak-ATG-Brn3a*-GSG-P2A-GGSGG-Runx1*-IRES-DsRed | CAG-Ngn2-DsRed (#576); BamHI, PmeI | pUCderivate-Brn3a-Runx1 (#1058); BglII, EcoRV, PvuI |
| pUCderivate-Brn3a-Ngn1 (#1065) | Kozak-ATG-Brn3a*-GSG-P2A-GSG-Myc-GGSGG-Ngn1* | gene synthesis, ATG:biosynthetics | |
| pUCderivate-Brn3a-Ngn2 (#1066) | Kozak-ATG-Brn3a*-GSG-P2A-GSG-Flag-GGSGG-Ngn2* | gene synthesis, ATG:biosynthetics | |
| CAG-Brn3a-Runx1-GFP (#1067) | CAG-Kozak-ATG-Brn3a*-GSG-P2A-GGSGG-Runx1*-IRES-GFP | CAG-Brn3a-GFP (#1056); BstXI, SfiI | CAG-Brn3a-Runx1-DsRed (#1064); BstXI, SfiI |
| CAG-Brn3a-Ngn1-DsRed (#1068) | CAG-Kozak-ATG-Brn3a*-GSG-P2A-GSG-Myc-GGSGG-Ngn1*-IRES-DsRed | CAG-Ngn2-DsRed (#576); BamHI, PmeI | pUCderivate-Brn3a-Ngn1 (#1065); BglII, EcoRV |
| CAG-Brn3a-Ngn2-DsRed (#1069) | CAG-Kozak-ATG-Brn3a*-GSG-P2A-GSG-Flag-GGSGG-Ngn2*-IRES-DsRed | CAG-Ngn2-DsRed (#576); BamHI, PmeI | pUCderivate-Brn3a-Ngn1 (#1065); BglII, EcoRV |
| pUCderivate-Prdm12 (#1080) | Kozak-ATG-Prdm12* | gene synthesis, ATG:biosynthetics | |
| CAG-Prdm12-DsRed (#1082) | CAG-Kozak-ATG-Prdm12*-IRES-DsRed | CAG-Ngn2-DsRed (#576); BamHI, PmeI (blunt) | pUCderivate-Prdm12 (#1080); EcoRV (blunt) |

*codon-optimized
